# AA_stat: intelligent profiling of *in vivo* and *in vitro* modifications from open search results

**DOI:** 10.1101/2020.09.07.286161

**Authors:** Lev I. Levitsky, Julia A. Bubis, Mikhail V. Gorshkov, Irina A. Tarasova

## Abstract

Characterization of post-translational modifications is among the most challenging tasks in tandem mass spectrometry-based proteomics which has yet to find an efficient solution. The ultra-tolerant (open) database search attempts to meet this challenge. However, interpretation of the mass shifts observed in open search still requires an effective and automated solution. We have previously introduced the AA_stat tool for analysis of amino acid frequencies at different mass shifts and generation of hypotheses on unaccounted *in vitro* modifications. Here, we report on the new version of AA_stat, which now complements amino acid frequency statistics with a number of new features: (1) MS/MS-based localization of mass shifts and localization scoring, including shifts which are the sum of modifications; (2) inferring fixed modifications to increase method sensitivity; (3) inferring monoisotopic peak assignment errors and variable modifications based on abundant mass shift localizations to increase the yield of closed search; (4) new mass calibration algorithm to account for partial systematic shifts; (5) interactive integration of all results and a rated list of possible mass shift interpretations. With these options, we improve interpretation of open search results and demonstrate the utility of AA_stat for profiling of abundant and rare amino acid modifications. AA_stat is implemented in Python as an open-source command-line tool available at https://github.com/SimpleNumber/aa_stat.

## INTRODUCTION

Bottom-up proteomics is widely applied for analysis of post-translational modifications (PTMs). Two main strategies are currently available: targeted PTM identification and the so-called unrestricted PTM search, the former being the most popular.[1–4] Targeted methods rely on *a priori* knowledge of modifications of interest for a proteomic sample, and are unable to characterize them in a blind manner. The latter is important for PTMs which have many types of functional groups, such as O-glycans.[5] Unrestricted search for PTMs can also facilitate studying the crosstalk between PTMs, which typically remains hidden in targeted searches. In addition, blind scanning for chemical artefacts associated with sample preparation can improve the results of subsequent peptide and protein identification and quantitation.

The concept of unrestricted PTM search appeared in the early 2000s.[6] However, the absence of powerful bioinformatics tools capable of handling vast search spaces within reasonable running time restricted implementation of the strategy. MS-Alignment was among the first algorithms proposed for unrestricted PTM search.[7] Around the same time TwinPeaks [8], SeMoP [9], OpenSea [10], SPIDER [11] and a number of other search tools [12–15] were introduced. Large-scale PTM profiling typically implied some restrictions of the search space due to processing time issues. As a result, detection of modifications of interest is usually only partially attainable.

This situation has started to change with wider adoption of ultra-tolerant search [16] facilitated by the appearance of the ultrafast search engine MSFragger.[17] The ultra-tolerant search uses a loose precursor mass tolerance (±500 Da and even more) for peptide identification. Here, we call this approach “open search”, while conventional, narrow-tolerance search is called “closed search”. Peptide-spectrum matches (PSMs) identified with the open search can be grouped by mass shift values that presumably correspond to either lost or attached functional groups. Recently, other engines, OpenPFind [18] and TagGraph [19], were proposed as fast alternatives to MSFragger for open search.

Introduction of new powerful tools for ultra-fast peptide identification using the open search strategy calls for comprehensive algorithms for postsearch validation and interpretation of detected mass shifts. Few such tools are available, and their functionality is typically restricted to mass shift clustering followed by filtering, validation and mass shift localization at a particular amino acid residue (PTMiner [20], DeltaMass [21]). Interpretation of the mass shifts is typically based on annotations in the Unimod database.[22] Recently introduced PTM-Shepherd aims at addressing a lack of metrics necessary for proper determination of identities and origins of modifications discovered with open search.[23] However, mass shift interpretation is still challenging when using any of the existing tools for post-processing of open search results. Indeed, many of the detected mass shifts either do not match known modifications, or cannot be interpreted as unaccounted missed cleavages, semi-tryptic digestion products and amino acid substitutions. Also, a lack of comprehensive and interactive visualization of the post-processed open search results further complicates generating hypotheses regarding the origin of mass shifts. Besides, there is still a need for established protocols to navigate discovery and annotation of rare post-translational modifications.

Recently, we introduced AA_stat, an open-source software for analysis of ultra-tolerant search results based on amino acid frequency calculation.[24] In this work, the AA_stat algorithm was significantly upgraded by adding MS/MS-based localization and implementing a number of features aimed at improving the method sensitivity and interpretation of the detected mass shifts. We demonstrate the utility of the updated AA_stat software for large-scale profiling of PTMs, as well as control of side products of complex sample preparation, PTM enrichment, labeling and solid phase peptide synthesis.

## EXPERIMENTAL SECTION

Publicly available proteomic data were used for the AA_stat evaluation.[25–32] Detailed annotation of these datasets is provided in Supporting Information (**S-Table 1**). Their selection was aimed at having a diverse number of modified amino acid residues per peptide. Specifically, the analyzed samples included enrichments of S/T phosphorylation, N-glycosylation, O-glycosylation, K-acetylation, as well as stable isotope labeling by amino acids in cell culture (SILAC)[33] and Tandem Mass Tags (TMT).[34] To estimate the localization accuracy, a synthetic dataset with known modifications was used.[35]

MSFragger v2.3 was used for both open and closed searches. Parameters for the open searches were default ones, except for the precursor and fragment ion mass tolerances and fragment ion types (**S-Table 2)**. SwissProt human canonical sequence database was used, retrieved on March 21, 2019. For the synthetic dataset, the database of synthesized sequences was appended to the human protein database. Decoy sequences were generated in reverse mode using Pyteomics.[36,37] Precursor and fragment ion mass tolerances, fragment ion types for subsequent closed searches were set as listed in **S-Table 3**. Isotope error correction, fixed and variable modifications for closed searches were set according to annotations (**S-Table 1**) or accounting for AA_stat recommendations (**S-Table 4**). Post-processing for closed searches was performed using Scavager.[38] Results for the open searches processed using MSFragger and AA_stat, as well as peptide tables produced by Scavager for closed searches, are available at https://gorshkovlab.github.io/aa_stat_reports/. Gene ontology (GO) enrichments were analyzed using STRING [39] (https://string-db.org/). A step-by-step description of the analysis of proline hydroxylation in proteins is provided in **Appendix A**(**Supporting Information**).

## RESULTS AND DISCUSSION

### Overview of AA_stat tool

AA_stat is an open-source command-line tool that processes the results of open search and produces a list of identified mass shifts that can be attributed to modifications. Its original version was described in detail in a previous work.[24] Briefly, AA_stat calculates and visualizes amino acid statistics within groups of peptides with shifted masses. This statistics allows to derive an initial hypothesis on possible modification sites. In the latest version, this statistics is complemented with MS/MS-based localization. In the course of operation, the AA_stat software performs the following steps (see **Figure 1**; recent additions to the algorithm are described in detail below): (1) reads and preprocesses the results of the open search (see *Mass calibration* below); (2) calculates a histogram of mass shifts observed in all considered PSMs; (3) applies smoothing to the histogram, followed by Gauss fitting on remaining local maxima; (4) applies group-specific FDR filtering to the mass shifts that passed the Gauss filter; (5) assigns a reference mass shift (see *Selection of reference mass shift* below) and calculates amino acid frequencies for peptides at each mass shift. These normalized frequencies indicate if a certain mass shift is “enriched” in a particular amino acid, prompting a hypothesis that peptides containing the given amino acid are *in vitro* or post-translationally modified, resulting in the observed mass shift; (6) performs (optionally) MS/MS-based localization of observed mass shifts. Here, possible localization sites are derived from observed amino acid frequencies, the Unimod database, and possible interpretation of mass shifts as combinations or as isotopic counterparts of other mass shifts (see *Localization of modification sites* below); (7) based on the results from (5) and (6), recommends a set of fixed modifications (for more sensitive open search) and variable modifications, as well as monoisotopic peak assignment error (for possible closed searches, or as a quick way to characterize the dataset) (see *Selection of fixed and variable modifications* below); (8) based on the results from (5) and (6), compiles a list of interpretations for all observed mass shifts, heuristically sorted by feasibility; (9) summarizes all results in an interactive HTML report, presenting the modification profile of the dataset; and (10) optionally, repeats open search using MSFragger, enabling fixed modifications recommended at step (7) (AA_search mode).

**Figure 1.**
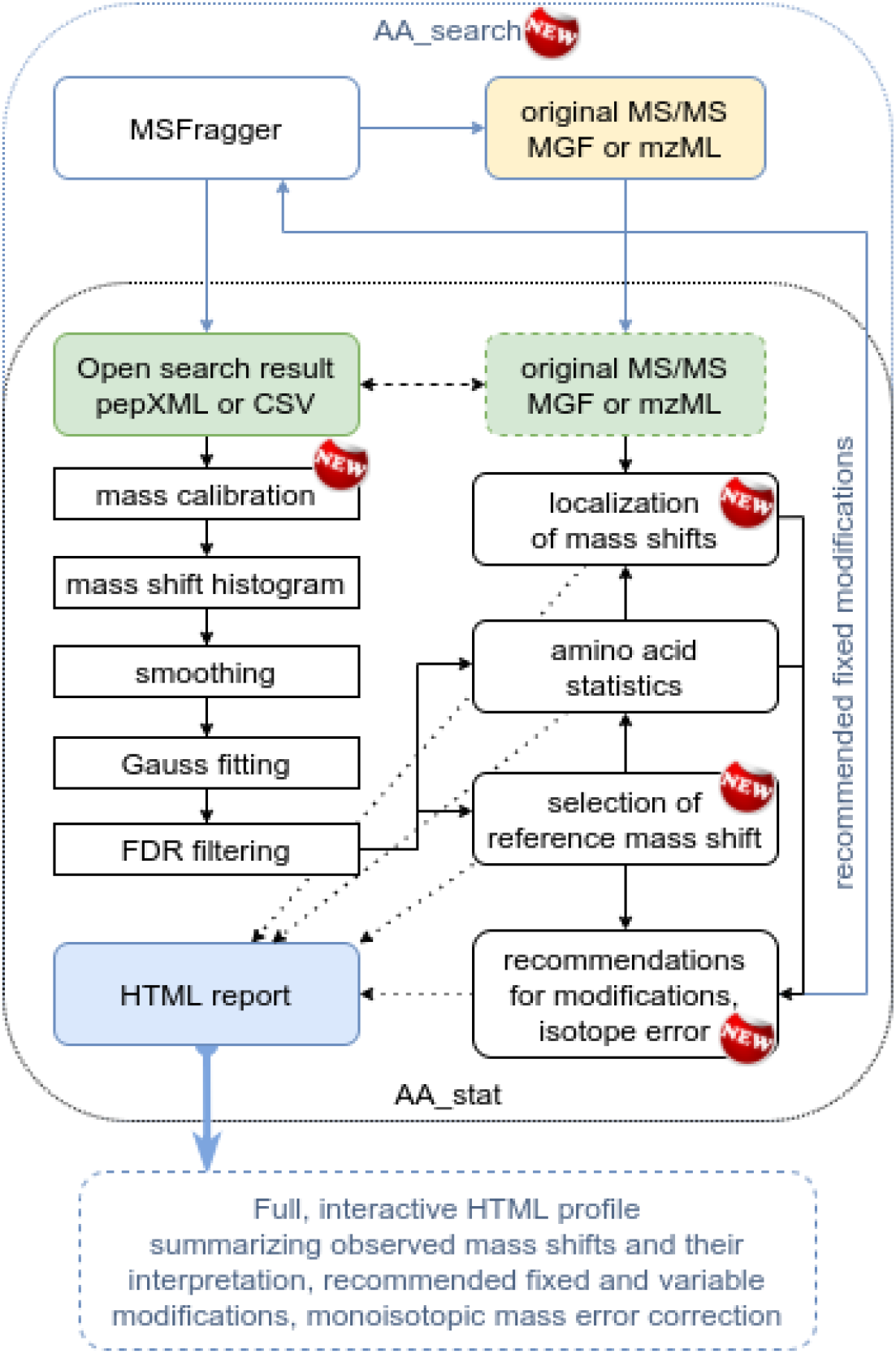
Overview of AA_stat and AA_search algorithms. AA_stat starts with the open search results (pepXML or CSV), then groups PSMs by mass shift values. Next, mass calibration, smoothing, Gauss fitting and group-specific FDR filtering are performed, followed by calculation of normalized amino acid frequencies. After composing the amino acid candidate list, AA_stat localizes mass shifts (using mzML or MGF files), concludes on probable fixed and variable modifications and produces a final HTML report. AA_search is a wrapper that runs an open search with MSFragger, followed by post-processing of the results with AA_stat. Performing in a cycle, AA_search can then apply fixed modifications recommended by AA_stat and repeat the open search for optimal results.

### Selection of reference mass shift

In the first version of AA_stat, the zero mass shift (containing unmodified peptides and/or peptides with fixed modifications) was always used for normalization of amino acid occurrence frequencies (*reference mass shift*). The latter restricted applicability of the tool to label-free and TMT-based proteomics workflows. For example, in case of metabolic labeling, zero mass shift can rarely be observed in the open search results and consequently provides poor normalization as a reference. The updated AA_stat dynamically selects the reference mass shift using the following algorithm. First, AA_stat determines if the zero mass shift exists in the window of [-0.05 Da, 0.05 Da]. Then, if such a mass shift is found and it contains more peptides than 5% of the most abundant mass shift, it becomes the reference mass shift. Otherwise, the most abundant mass shift is chosen as the reference. If the zero mass shift is found, it is utilized for mass calibration for mass calibration to correct possible systematic mass measurement errors.

### Mass calibration

During the preprocessing step when reading individual input files, in case the zero mass shift contains enough peptides (500 peptides with 2% FDR within ±0.05 Da), it is used to estimate and correct for systematic mass measurement errors. First of all, the filtered peptides are considered to determine if there are any sudden shifts in mass measurement errors in the course of the experiment. To achieve this, the two-dimensional array of peptide retention times and mass errors is scaled and then supplied to the DBSCAN clustering algorithm [40,41] implemented in scikit-learn library.[42] Obtained clusters are then filtered by size and duration. If more than one significant cluster is identified, the whole set of identifications is divided by retention times and each slice is assigned to one of the clusters. Mass errors of unmodified peptides in each of the clusters are fit to a Gaussian function. The derived mean of the Gaussian is the calculated systematic mass shift for the cluster, and all identifications in the assigned retention time range are shifted to correct this systematic error.

Instead of absolute mass units, AA_stat can perform clustering and Gauss fitting using relative mass units or “frequency units”. The latter is the default and can be used for data obtained using Orbitrap mass analyzers.[43] In this mode, measured masses are converted back to frequencies (according to the mass calibration equation) prior to cluster analysis and Gauss fitting. **S-Figure 1** shows an example use case for dataset #3 (see **S-Table 1**), wherein a partial shift in the middle of one experimental run is corrected with clustering.

### Localization of modification sites

AA_stat localizes identified mass shifts and their superpositions using MS/MS spectra. The localization procedure is outlined below.

First, a list of possible amino acid localizations for each mass shift is formed. There are four sources of amino acid candidates (eligible sites) for a specific mass shift: (1) all amino acids (including N- and C-terminus) from Unimod database that correspond to the mass shift with a given mass tolerance; (2) amino acids whose normalized frequency [24] exceeds a given threshold (1.5 by default); (3) if a mass shift can be a ^13^C isotope within a given mass tolerance, all candidates for the monoisotopic mass shift are considered; (4) if a mass shift equals the sum of two other observed mass shifts, simultaneous localization is attempted using their respective localization sites. Alternatively, AA_stat can be configured to consider all amino acids as possible localization sites.

At the next step, the isoforms for each peptide are generated according to the amino acid candidate list. If the considered mass shift equals the sum of two other observed mass shifts within a given tolerance, isoforms for the combination of modifications are also generated.

Next, the theoretical spectrum for each peptide isoform is matched against the experimental spectrum using a simplified renormalized Hyperscore scoring function introduced earlier.[44] If there is no clear winner among the isoforms, the mass shift is treated as “non-localized”. If a modification is localized at a terminal residue, then it is also considered to be localized at the corresponding peptide terminus. Localization score is calculated as the difference between the top isoform score and the second best one, divided by the top score.

Localization results are summarized in CSV tables, containing top isoforms, experimental spectrum IDs and localization scores for each identified peptide at the considered mass shift. Localization statistics for specific mass shifts along with normalized amino acid frequencies and peptide percentages are shown in self-titled figures.

### Selection of fixed and variable modifications

The new version of AA_stat infers the amino acids that are highly or completely modified in a dataset. Although fixed modifications introduced during sample preparation are usually known to data holders, there are several reasons why such an option was implemented. First, increasing the abundance of the zero mass shift by applying fixed modifications improves the sensitivity of the method and the accuracy of amino acid statistics calculated by AA_stat (dataset #6). Second, we found that application of multi-step sample preparation with peptide enrichment can result in peptide fractions with unexpected complete modification (methionine oxidation, datasets #6 and #11 in this study). Thus, this option provides more information on the consequences of the applied steps of complex sample preparation [45] and is of potential interest for characterization of efficiency and side products of new techniques for enrichment and labeling. Third, from our experience, simply wrong, or incomplete annotation of sample preparation unfortunately occurs.[24] If the alkylating agent is erroneously specified, the option of inferring fixed modifications from search results will automatically correct the mistake without additional efforts from the researcher.

Details of the algorithm for selection of fixed modifications are shown in **S-Figure 2A**. Briefly, the algorithm selects the optimal modification to compensate the non-zero reference mass shift or to increase the low peptide percentages observed at the zero mass shift. AA_search routine has an option that implements the recommended fixed modifications and runs another round of open search automatically. This is repeated until no more fixed modifications are needed. This typically takes from one to four steps, depending on the complexity of the data. Note that the fixed modifications suggested by AA_stat do not necessarily correspond to sample preparation; it is also not always optimal to set them as fixed in the subsequent closed search, although, it can be appropriate, if the amino acid modification rate is close to 100%. Recommended fixed modifications are meant as a suggestion for the open search to improve identification rate and the accuracy of AA_stat results.

Variable modifications are recommended to summarize the most abundant mass shift localizations determined by AA_stat (top 5 by default). With this option, AA_stat aims at producing the most effective list of variable modifications for the follow-up closed search. Recommendation is done solely on the basis of successful MS/MS localizations, preferring the mass shifts and modification sites with the highest absolute localization count. Already recommended fixed modifications (and already enabled variable modifications, if any) are excluded. The user can also instruct AA_stat to not recommend multiple modifications of the same residue. Additionally, a recommendation is made to allow for the possibility of isotope mass error; it is also accounted for in the selection of mass shifts as variable modifications. Exact procedure is outlined in **S-Figure 2B**.

### Accessing the results of mass shift profiling and interpretation

The easiest way to obtain the open search and postsearch processing results is running MSFragger and AA_stat in a single command using AA_search (**Figure 1**). The results of mass shift profiling and interpretation are summarized in the interactive HTML report (**Figure 2**) (examples can be found at https://gorshkovlab.github.io/aa_stat_reports/). At the top of the report, the fixed and variable modifications recommended by AA_stat are shown (**Figure 2A**). Variable modifications correspond to the most abundant mass shift localizations in the dataset and can be used to maximize the yield of closed searches. Fixed modifications are recommended if unmodified peptides are not identified or a particular amino acid is not found in them. These can appear due to unspecified or incorrectly specified labeling, alkylating agents, or due to absence of unmodified peptides (to compensate a loss in method’s sensitivity). Below, the table summarizing all identified mass shifts, number of peptides per mass shift, statistics of amino acid occurrence and mass shift interpretation is provided (**Figure 2B**). Below the table, there is a bar plot visualizing all detected mass shifts and relative numbers of peptides in each (**Figure 2C**). Mass shift values are interactively linked to bar plots visualizing amino acid occurrence relative to the reference mass shift, percentage of peptides containing the particular amino acid residue and counts of mass shift localizations at each particular amino acid (the plot appears below the statistics table, replacing the mass shift histogram) (**Figure 2D**). Clicking on the number of peptides at the given mass shift (“# peptides in bin” column) opens the table containing the modified peptide sequences, modification sites, scores of site localization and spectrum identifiers (**Figure 2D**). Mass shift interpretation (“Possible interpretations” column, **Figure 2B**) contains: (1) interactive links to the Unimod database, if the mass shift has Unimod annotation, (2) sums of mass shifts, if a given mass shift can be interpreted as a combination of two others, (3) reference to the monoisotopic mass shift, if a given mass shift can be interpreted as an isotope of another, (4) possible explanations for artefact mass shifts.

**Figure 2.**
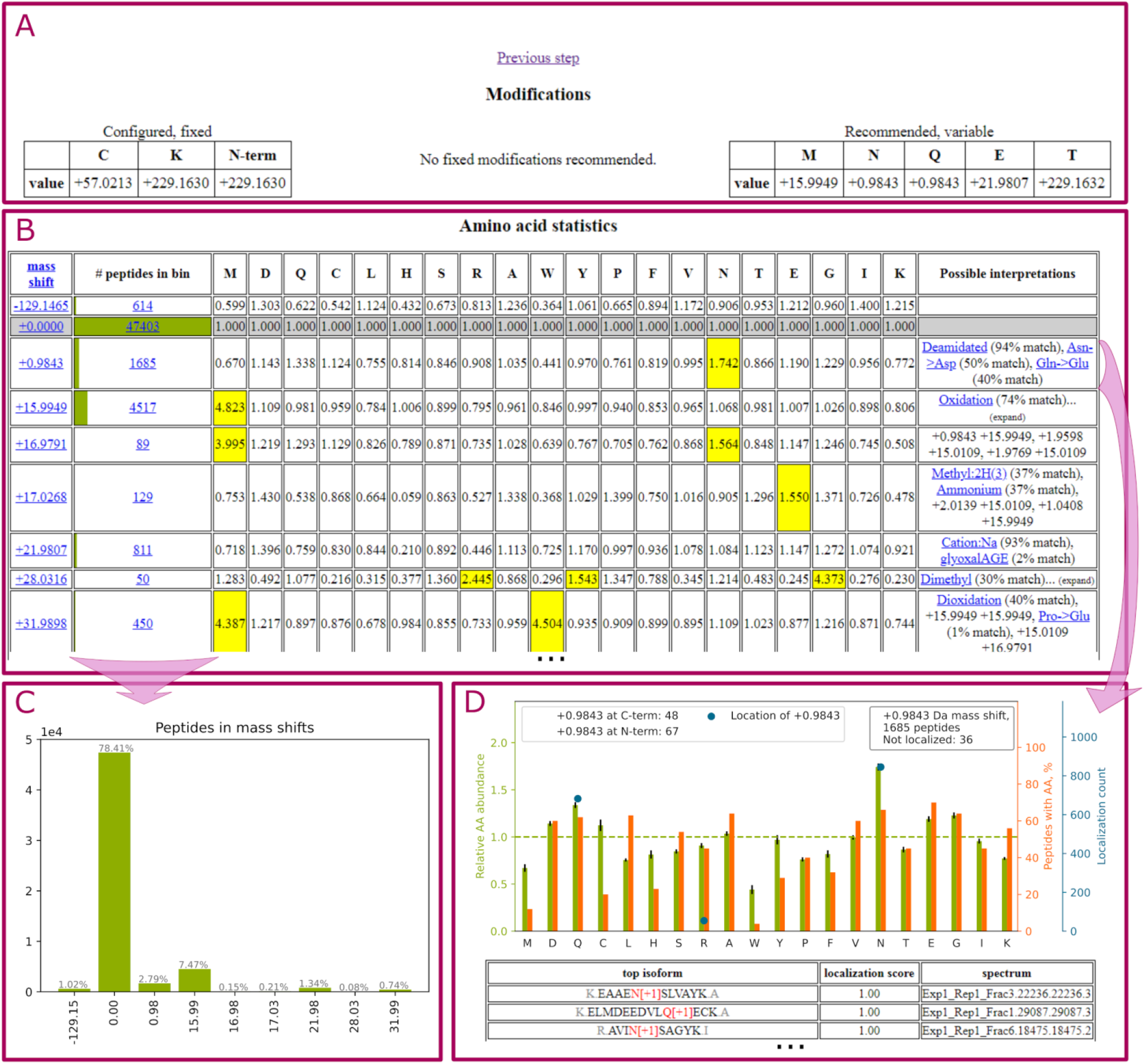
AA_stat report summarizes the results of open-search-based modification profiling and its interpretation. **A:** the top section summarizes detected and recommended modifications. “Configured, fixed” (left) - fixed modifications detected in search parameters by AA_stat (i.e. already enabled); “Recommended modifications” (center): if present, shows AA_stat recommendation for fixed modifications; ‘’Recommended, variable” (right): AA_stat recommendation for variable modifications in closed search. This also summarizes top localizations at a glance. **B:** Main statistics table, listing all mass shifts, number of peptides bearing each mass shift, and normalized occurrence frequencies of all amino acids. Highlighted in yellow are enriched amino acids. The “Possible interpretations” column lists possible interpretations of the mass shift: Unimod URLs, representation as isotopes or combinations of other observed mass shifts, or artefacts. **C:** Mass shift histogram, showing all observed mass shifts and normalized numbers of peptides identified with these mass shifts. **D:** When clicking on a mass shift, the main histogram is replaced with a figure characterizing the selected mass shift. It shows normalized amino acid occurrence frequencies (light green bars), percentage of peptides containing each amino acid (orange bars), and successful localization counts for each amino acid (blue markers). In case a mass shift can be interpreted as a sum of two others (in one or more ways), localization is done based on all possible interpretations of the mass shift and shown with different markers.

Along with the HTML report AA_stat generates figures with zero mass shift clustering results for each file, PDF file with Gauss fit results for each mass shift candidate, tables and figures for each mass shift summarizing amino acids statistics, localization results and peptide lists. Description of all generated files can be found on the GitHub page of the project (https://github.com/SimpleNumber/aa_stat).

### Assessment of AA_stat performance using enrichment and labeling data

The enrichment and labeling data represent a convenient resource for estimation of the accuracy and sensitivity of the developed algorithms: modifications introduced during sample preparation are well known, and they can model a complex case when multiple modifications in a peptide should be resolved. Thus, a competent tool should be able to recognize from the open search results the complex combination of intentionally introduced modifications. **S-Table 4** summarizes AA_stat output for 16 datasets: top mass shift localizations by abundance are shown, suggesting users to consider probable fixed or variable modifications. AA_stat has expectedly reported the carbamidomethylation of Cys, TMT and dimethyl labels as fixed modifications, while SILAC labels, enriched phosphorylation, N- and O-glycosylation, post-translational and artificial acetylation, methionine oxidation and some other artificial modifications are identified as probable variable modifications. Correction of the monoisotopic peak assignment errors, occurring frequently in proteomic experiments,[46] was recommended for 3 of 16 datasets.

After comparison of the modifications annotated from the respective studies and the AA_stat-retrieved abundant mass shift localizations, (**S-Table 4)** unexpected consequences of complex sample preparation can be noted. Met oxidation is always believed to be variable modification and used as such. However, AA_stat has reported “fixed” oxidation at Met residues for datasets #6 and #11 (https://gorshkovlab.github.io/aa_stat_reports/). This means that Met oxidation level can be very high in these particular datasets. The closed searches for #6 and #11 with variable methionine oxidation further supported this observation and demonstrated the Met oxidation rates of 100 and 99.2%, respectively (**S-Figure 3**). This suggests that complex sample preparation including peptide enrichment can result in peptide fractions with by-products, i.e. containing “enrichment” of the unexpected modifications. The fixed modifications suggested by AA_stat can, thus, provide useful knowledge about the “side effects” of sample preparation.

The dataset with R-methylation enrichment (#12 in **S-Table 4**) contains ~500 peptides with 57.02 Da mass shift, however, fixed Cys carbamidomethylation was not suggested. Besides, although a mass shift of 14.02 Da corresponding to R-methylation was detected in #12, it was of low abundance and was not among the top seven mass shift localizations. Closed searches performed for #12 revealed peptide identifications containing less than 0.5% of peptides with carbamidomethylation of Cys and less than 0.5% of peptides with methylation or dimethylation of Arg (~200 of 42,000 peptides). This can again suggest the specific consequences of complex sample preparation and enrichment resulting together in a significant loss of Cys-containing peptides. Low number of R-methylated peptides can be due to consideration of a small part of the whole dataset. However, insufficient performance of R-methylation enrichment can also be the case.

Dataset #16 (CPTAC colon cancer, sample 01) was found to contain approximately 7,000 overalkylated peptides: mass shifts of +57.02 and +114.04 Da were predominantly localized at the N-terminus.

For dataset #6 enriched with N-glycosylation, AA_stat assigned the most abundant mass shift (+0.984 Da at Asn) as a fixed modification to compensate for the absence of unmodified peptides at the zero mass shift. Since no known modifications were pre-set in the open search, this step was required for proper calculation of amino acid statistics and was automatically applied to increase the sensitivity of the method. This artificial step can be recognized by the appearance of −0.98 Da at Asn in the list of probable variable modifications. The result should be treated as a recommendation of variable modification of +0.984 Da at Asn.

In datasets #13-15, +128 Da at the N-terminus was among the top seven mass shifts. It corresponds to extra lysine at peptide terminus. This mass shift dissipates when a larger number of missed cleavages is accounted for in the open search (**S-Figure 4**). For the closed searches, it can also be considered as a recommendation to increase the number of missed cleavages.

Some search engines restrict the search space by limiting the number of variable modifications per peptide. Based on the results in the AA_stat HTML report, a grounded decision on the value for the parameter can be made. Detailed instructions for such a decision are provided using AA_stat results for dataset #7 (**S-Figure 5**). This subset combines abundant post-translational Lys acetylation (42.01 Da), metabolic labeling of Lys and Arg residues (8.01 and 10.01 Da) and methionine oxidation (15.99 Da). Specifying one missed cleavage suggests the presence of up to two modified lysines per peptide. Thus, taking into account the percentage of peptides bearing together methionine oxidation, SILAC labels, and acetylation, allowing recommendation of at least three sites per peptide.

Thus, AA_stat allows controlling the efficiency and side reactions of complex sample preparation. Variable modifications recommended by AA_stat can recover up to 15% additional peptide identifications in the closed searches (#12 in **S-Figure 6B**).

### AA_stat demonstrates potential for uncovering rare modifications

One of the common strategies for mass-spectrometry-based discovery of rare post-translational modifications is the analysis of big public data followed by validation of the collected rare events for significance and biological relevance.[47] We tested AA_stat for the ability to accumulate peptides with rare PTMs when analyzing a subset of data and demonstrated its potential for uncovering rare PTMs. Oxidation of proline was used as an example. Proline oxidation is common in collagens, but rarely observed in other proteins. Based on the AA_stat reports, we selected from the CPTAC dataset (one colon cancer sample, #16 in **S-Table 4**) the mass shift of 15.99 Da containing, among the others, 70 peptides with localization of hydroxyl group at Pro. These peptides predominantly originated from collagens and collagen-like proteins, a well-characterized protein family which needs hydroxylation of prolines for correct folding.[48] GO analysis of the corresponding proteins revealed enrichment of collagen trimer and endoplasmic reticulum (ER) where hydroxylation takes place (**S-Table 5**, **S-Figure 7**). Localization of hydroxyl groups at particular protein sites was additionally verified against the Uniprot database. 26 of 70 localizations reported by AA_stat are at sites listed in Uniprot (**S-Table 6**). Consensus-motif recognition demonstrated the presence of XpG (any amino acid - hydroxyproline - glycine) motif around localization of hydroxyproline for 53 of 70 cases. XpG is a well-described conserved amino acid motif for the prolyl 4-hydroxylases, the enzymes catalyzing modification of collagens in ER.[49]

Next, we ran AA_stat across a hundred of colon cancer samples available from CPTAC and then again retrieved from AA_stat results the PSMs with hydroxyl groups localized at prolines. Besides collagens (with and without XpG motif), we observed six cases of proline oxidation in peptides with a conserved amino acid pattern, F(Y or F)ApWCGHC(K or Q), that corresponds to protein disulfide-isomerases (PDIA1, PDIA3, PDIA4, PDIA5, and PDIA6) and thioredoxin domain-containing protein (TXNDC5) (spectra are shown in **S-Figure 8**). These modification sites are not annotated in the PTM databases (Uniprot, iPTMnet) or literature in the context of colon cancer proteomics and can be potentially new. The only observation of the hydroxylation of prolines in the protein disulfide isomerases has been reported earlier for HeLa cells.[47] Also, one case of unannotated proline oxidation in peptides from human histones was observed (spectrum is shown in **S-Figure 9**). The question of validation of potentially new histone hydroxylation remains open. Since other methods capable of providing complementary experimental evidence are absent, the approach to proving the new modification sites can be seen in cross-validation of the results by consideration of larger sample cohorts. Thus, starting with one deep proteome we were capable of reproducing common proline hydroxylation in collagens, while by moving to a hundred samples we reproduced the proline hydroxylation on both collagens and protein disulfide isomerases and found potentially new hydroxyprolines in human histones. Therefore, while approximately 50 peptides are enough for initial detection of a mass shift with AA_stat (e.g. hydroxylation of methionine), larger datasets allow discerning less common modification sites (e.g. proline), and with growing data size, rare and unique PTMs corresponding to this same mass shift. Therefore, we conclude that the AA_stat-based open search analysis workflow is applicable for discovery and validation of modifications rarely observed in mass spectrometry data, including rare post-translational modifications. We provide a step-by-step description of analysis implemented for hydroxyproline characterization (**Appendix A** in Supporting Information).

### Limitations, pitfalls and artefacts of the open search strategies decrease sensitivity of modification profiling

The standard enrichment and labeling data used as model datasets helped us to assess the limitations of the open-search strategies, aimed at uncovering multiple modifications of a peptide sequence (**S-Table 1**). **S-Figure 10** demonstrates that phosphorylation enrichment followed by TMT labeling (dataset #1) resulted in a peptide set in which more than 90% of peptides bear three or more modifications. The presence of two or more modifications in the peptide sequence shifts the fragment masses observed in tandem spectra. If modification sites are distantly located from each other, the chance of matching the MS/MS spectrum of a modified peptide to the theoretical spectrum of its unmodified isoform significantly decreases. Particularly, less than 10% of tandem spectra in such “over-modified” datasets can result in successful identifications in open search (**S-Table 2**). Another example are phospho-enriched SILAC-labeled datasets (#9 and #10 in **S-Table 1**) suffering the same limitations. Two modifications located at N-terminus and C-terminus prevent identification of the peptide in open search. One more complicating issue for enriched samples is the total number of peptides: a few hundred peptides distributed across all mass shifts are not enough to pass the AA_stat filters (Gaussian fitting and group-specific FDR).

The above factors together affect method performance. For example, cysteine carbamidomethylation in datasets #9 and #10 mostly occurs along with Ser or Thr phosphorylation and will not be observed as a separate mass shift if SILAC labels are not set as variable modifications (https://gorshkovlab.github.io/aa_stat_reports/). A similar issue is observed in datasets #4 and #5 with O-glycosylation enrichment, where light (~28Da) and heavy (~32Da) dimethyl labels are used. Although HexNAc (N-acetylglucosamine) with a mass shift of +203 Da localized at Ser and Thr is detected, dimethyl labels are not properly determined (**S-Table 4**) due to insufficient number of modified peptides matching the unmodified spectrum and passing Gaussian fit and FDR filtering (**S-Figures 10 and 6A**). A possible solution in these cases is the inclusion of known modifications (i.e. carbamidomethylation, labels and/or enriched PTM) in the open search, however, at the price of a significantly increased running time.

In the above examples, the modifications are known in advance; however, the same problem can occur with peptides bearing multiple chemical and/or post-translational modifications. When modifications are located in close proximity to each other, the peptide is more likely to be identified in open search. However, this will result in observing complex mass shifts, each representing a sum of modifications. The current version of AA_stat attempts interpretation of mass shift combinations (**Figure 3**). If any of the mass shifts composing the possible combination are predominantly localized as a sum of two other observed mass shifts, the presence of more than two modifications in the peptide sequence can be suggested. With a large number of detected mass shifts, an exponentially high number of combinations results in spurious mass coincidences and incorrect localizations.

**Figure 3.**
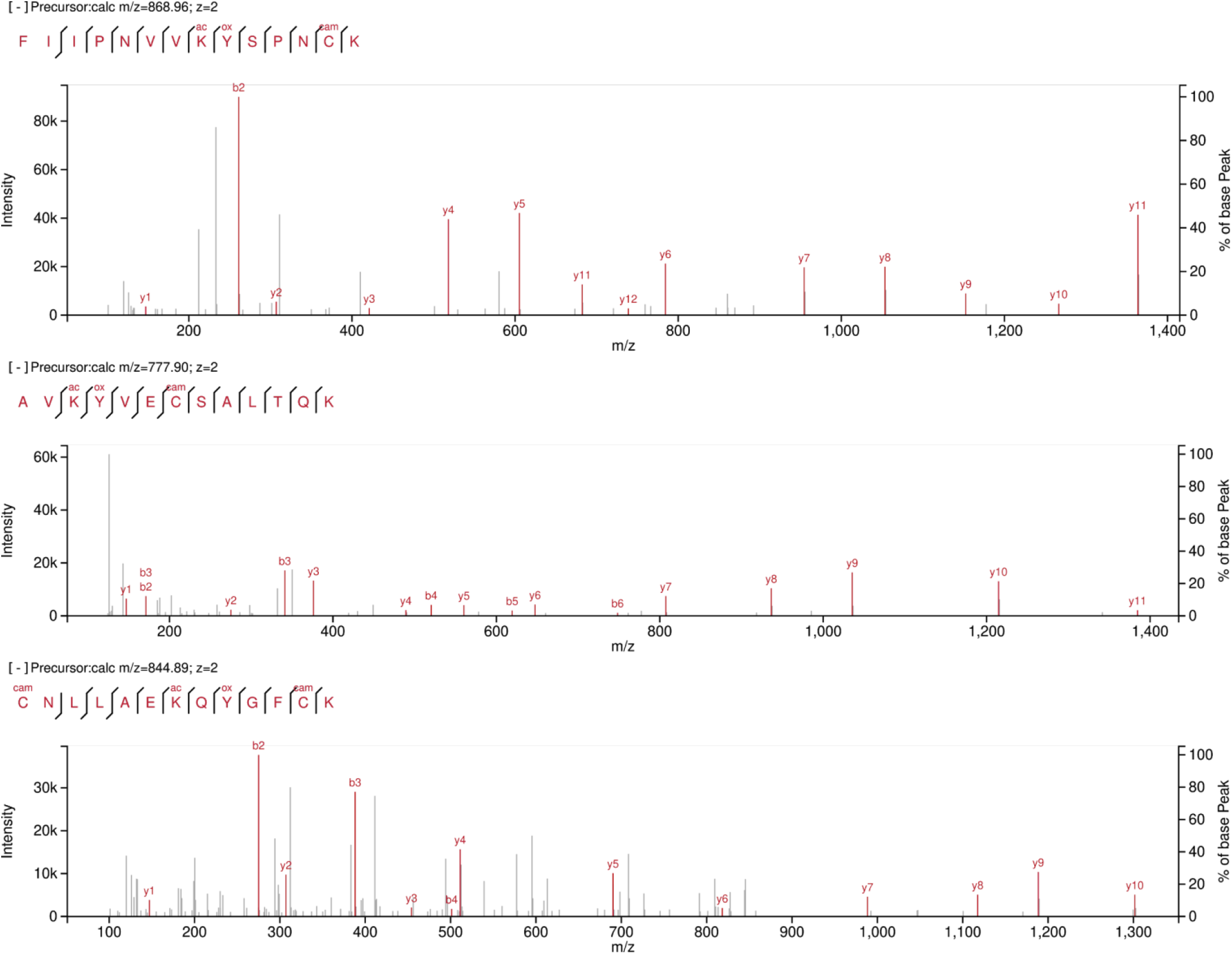
Mass spectra of multiply modified peptides with localization of two post-translational modifications using AA_stat (subset #7 in **S-Table 1**).

Many of the mass shifts that passed Gauss fitting and group-wise FDR filtering do not match known modifications, cannot be interpreted as a sum of other modifications, monoisotopic error, unaccounted missed cleavage, semi-tryptic digestion products and are not amino acid substitutions despite matching one within a given accuracy. For example, in dataset #16, mass shifts −128, −71 and −99 Da occur artificially. These mass shift match the masses of Lys-loss; Lys-loss + 57.02 and Arg-loss + 57.02 and are localized at the terminal amino acid residue that is lysine or arginine (AA_stat report #16 at https://gorshkovlab.github.io/aa_stat_reports/). In open search, the tolerance for precursor mass typically covers ±500.0 Da. Thus, KPEPTIDER and PEPTIDER are suitable for the same mass spectrum. The MSFragger engine scores KPEPTIDER higher (X!Tandem hyperscore relies on the number of the matched ion fragments and their intensities), if abundant b_1_ or y_n_ fragment ions are occasionally matched (**S-Figure 11**). As a result, a mass shift matching N-terminal amino acid loss can occur. Currently, AA_stat provides an interpretation of the possible artificial origin for such a mass shift.

Some uninterpretable mass shifts originate from technical instabilities of the instrument. For example, the first sample fraction in dataset #3 has a systematic mass shift from 55th to 80th minute of the gradient (**S-Figure 1A** in Supporting Information). If the calibration algorithm does not account for this effect, uninterpretable mass shifts are observed (0.028 Da, 15.99 + 0.028 Da in **S-Figure 1A**). The clustering-based calibration algorithm implemented in the new version of AA_stat accounts for such effects (**S-Figure 1B)**.

### Estimation of localization accuracy using synthetic peptides with different modifications

To evaluate the localization accuracy, we used a publicly available dataset obtained for the sample containing ~5000 modified synthetic peptides.[35] The open search results processed with AA_stat are provided at https://gorshkovlab.github.io/aa_stat_reports/. The accuracy of localization was defined as the percentage of correctly localized synthetic isoforms reported by AA_stat for a particular mass shift among the total number of peptides at this mass shift. This parameter varies from 75% to 95% depending on modification (**S-Figure 12**). A decreased localization accuracy (from 54% to 58%) was observed for mono-, di- and trimethylation. In the original study, synthetic peptides with these modifications were characterized by difficulties in solid phase synthesis and low ion fragmentation efficiency. Other studies reported on characteristic neutral losses for methylated peptides, as well as loss of other peptide backbone fragments.[50] In addition to the modified synthetic peptides annotated in the original study, open search reveals peptides differing by one amino acid, or corresponding to a loss of modification, or having other modifications (e.g. https://gorshkovlab.github.io/aa_stat_reports/synthetic/Phospho/report.html). Thus, we conclude that AA_stat demonstrates accurate localization and can be used to estimate: (1) purity of synthetic peptides (chromatographic purification may not be sufficient to remove all side products), and, (2) other modifications occurring during sample preparation and MS analysis (e.g. *in vitro* modifications, side chain losses, etc). These questions remain open for any synthetic dataset, however, open search analysis combined with AA_stat has a high potential for reconstructing the profile of side products in peptide synthesis.

### Comparison with other tools for post-open-search analysis

A number of software tools have been presented to date for open search interpretation and/or PTM localization.[8–12] Among them, we identified PTM-Shepherd [23] as the closest to the one described here since it reportedly produces a similar output and utilizes a similar approach to postsearch validation. However, PTM-Shepherd is part of FragPipe [17] and requires multiple prior steps of analysis. AA_stat, on the other hand, is a standalone command-line tool invoked in a single command after obtaining the open search results. Both PTM-Shepherd and AA_stat produce text tables with information on detected mass shifts and results of their localization, but AA_stat additionally plots summarizing statistics and visualizes all results in a single, interactive HTML report (**Figure 2**).

We compared the tools using the whole colon cancer data from CPTAC study S037 to assess the differences and complementarity in the results (**S-Figures 13-14**). On the CPTAC dataset, AA_stat reports 343 mass shifts. Of those, 59% coincide with mass shifts reported by PTM-Shepherd within 0.01 Da tolerance (**S-Figure 13A**). Considering top 100 mass shifts reported by both tools, the overlap increases to 64% (**S-Figure 13B**). Note that PTM-Shepherd reports top 500 mass shifts, while AA_stat does not have that limitation. The number of mass shifts reported by AA_stat is generally lower with default settings, due to different approaches to mass shift selection and FDR control. AA_stat employs group-specific filtering (at 2% by default) using “+1” correction [51] and frequency calculation at the peptide level. PTM-Shepherd operates at PSM level and does not perform group-specific filtering. As a result, most differences are expected on low-abundance mass shifts.

Next, we analyzed the peptides at mass shifts reported by both PTM-Shepherd and AA_stat. We compared the numbers of peptides per mass shift (**S-Figure 14**). Despite the above-mentioned differences in procedure, the abundances of reported mass shifts tend to agree, although less so for less abundant mass shifts.

Finally, we considered localization sites determined by PTM-Shepherd and AA_stat (**Figure 4**). For 40 most abundant mass shifts identified by both tools, localizations are largely in agreement. Major differences are observed for the mass shifts of 0.93, −0.92, −0.96, −1.93 and 1.92 Da, which have poor localization rates (0-38%) meaning the mass shifts were basically not localized by both PTM -Shepherd and AA_stat. Such unsuccessful localization can be attributed to probable artefacts of the search, data processing or acquisition; an unresolved sum of modifications is also possible. The mass shift of 17.02 Da matches a replacement of proton with ammonium ion. Unimod database reports Asp and Glu as preferable sites, and AA_stat demonstrates a better localization coincidence with Unimod database than PTM-Shepherd. The 22.98 Da mass shift is resolved by AA_stat as a probable isotope of +21.98 Da, which matches sodium adduction on Glu and Asp. Mass shift of - 128.09 Da matches Lys and Gln masses and can be explained by open search artefacts, at least when localized at peptide N-terminus (as stressed above in this paper). Similarly, −71.07 Da was interpreted by AA_stat as a sum of −128.09 and 57.02 Da. Current version of PTM-Shepherd lacks the ability to interpret the mass shifts as a combination of others. The mass shift of 186.03 Da has no match in Unimod database. It is preferably localized by AA_stat on Lys, while PTM-Shepherd determines preferable localization on Leu. We assume that 186.03 Da can be a sum of more than two modifications that was unresolved by both tools.

**Figure 4.**
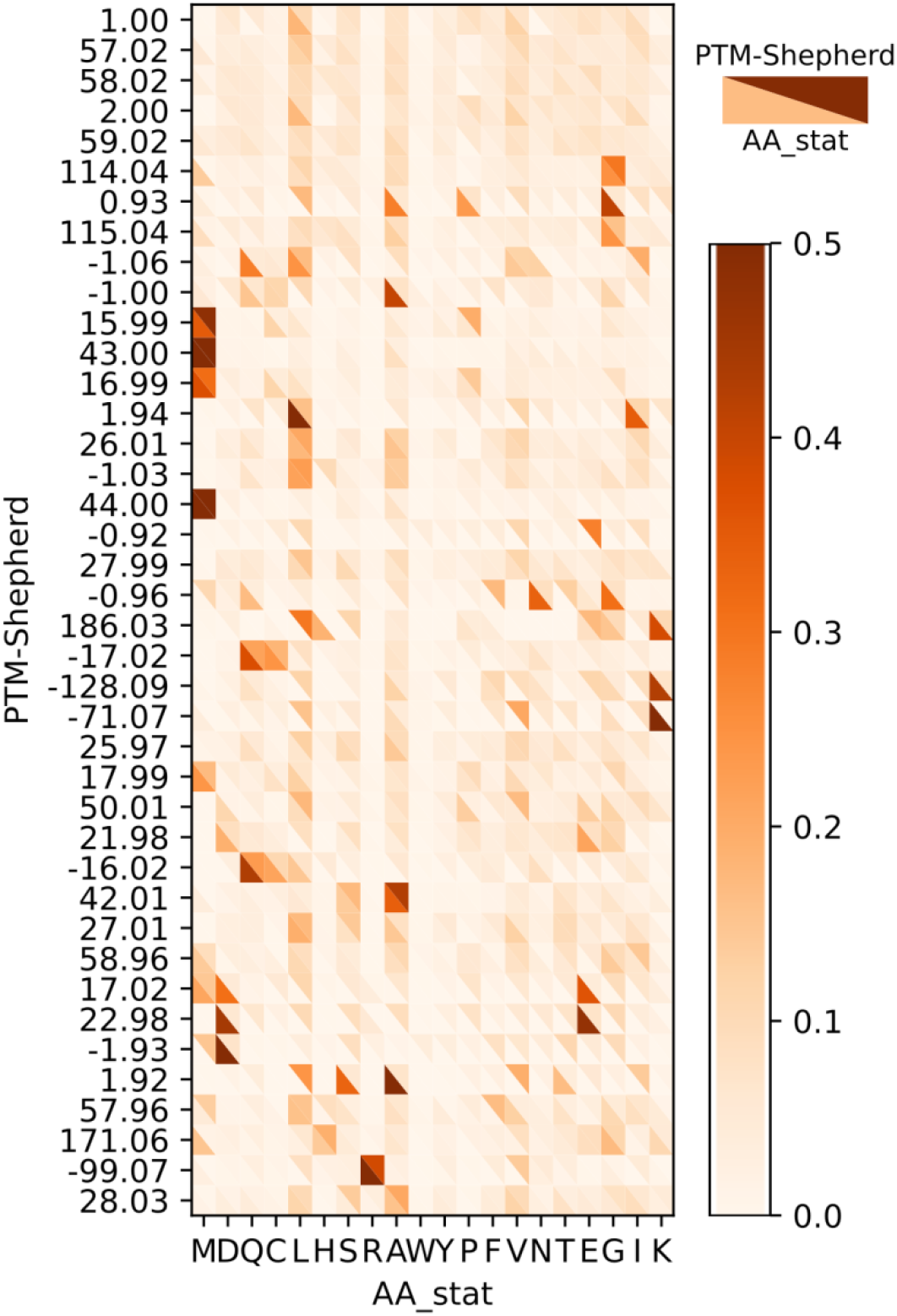
Comparison of AA_stat and PTM-Shepherd localization: top 40 most abundant overlapping mass shifts. Bottom-left triangles represent AA_stat results, top-right triangles -- PTM-Shepherd results. All values are normalized to the number of localized peptides (AA_stat) and PSMs (PTM-Shepherd). Mass shifts are sorted by peptide numbers as reported by AA_stat, with most abundant mass shifts at the top.

In general, there are significant differences between mass shifts reported by PTM-Shepherd and AA_stat, especially the less abundant ones. Each of the tools reports unique mass shifts due to different algorithmic solutions, therefore, the findings potentially complement each other. Among the mass shifts reported by both tools, there is a good agreement in localization and in the numbers of peptides for most of the overlapping mass shifts. The differences in mass shift localization can be attributed to: (1) variances in peptide populations determined for the mass shift; (2) PTM-shepherd can localize only a single modification per peptide, while AA_stat attempts localization of two modifications; (3) PTM-shepherd and AA_stat may use different sets of potential localization sites when localizing the same mass shifts.

## CONCLUSION

The AA_stat tool developed for post-processing of open search results was upgraded with addition of MS/MS-based localization of mass shifts and localization scoring, including shifts which are the sum of modifications. Interactive integration of post-processing results and a sorted list of possible mass shift interpretations implemented in the new AA_stat help to determine the origin of the mass shifts. The updated software accounts for systematic mass shifts in a part of the experimental run, has acquired an improved sensitivity for multiply modified peptides, as well as rare PTMs, recognizes monoisotopic peak assignment errors and infers most abundant mass shift localizations that can be further used to increase the yield of closed search. Generation of dataset-specific profiles of modifications allows accounting for unexpected modifications, and can be considered as a first step to blind data analysis without *a priori* knowledge of experimental conditions. These improvements to the AA_stat software make it a versatile and robust tool for comprehensive modification profiling.

## ASSOCIATED CONTENT

Supporting Information is available.

## ACKNOWLEDGEMENTS

Authors thank Dr. Mark Ivanov for helpful discussions on AA_stat development. Data processing was performed with the support from the 2013-2020 Program of Basic Research of the Russian State Academies of Sciences. Bioinformatic tool development was supported by Russian Foundation for Basic Research (# 18-29-13015).

## AUTHOR CONTRIBUTIONS

I.A.T. conceived the project, L.I.L. and J.A.B. developed the software and contributed equally, I.A.T., L.I.L. and J.A.B. performed analyses, all authors wrote the manuscript, M.V.G. and I.A.T. supervised the entire project.

## REFERENCES

[1] Falkner, J.A., Falkner, J.W., Yocum, A.K., Andrews, P.C., A spectral clustering approach to MS/MS identification of post-translational modifications. J. Proteome Res. 2008, 7, 4614–4622.

[2] Frank, A., Pevzner, P., PepNovo: De novo peptide sequencing via probabilistic network modeling. Anal. Chem. 2005, 77, 964–973.

[3] Frank, A., Tanner, S., Bafna, V., Pevzner, P., Peptide sequence tags for fast database search in mass-spectrometry. J. Proteome Res. 2005, 4, 1287–1295.

[4] Tharakan, R., Edwards, N., Graham, D.R.M., Data maximization by multipass analysis of protein mass spectra. Proteomics 2010, 10, 1160–1171.

[5] Xiao, H., Sun, F., Suttapitugsakul, S., Wu, R., Global and site-specific analysis of protein glycosylation in complex biological systems with Mass Spectrometry. Mass Spectrom. Rev. 2019, 38, 356–379.

[6] Tsur, D., Tanner, S., Zandi, E., Bafna, V., Pevzner, P.A., Identification of post-translational modifications by blind search of mass spectra. Nat. Biotechnol. 2005, 23, 1562–1567.

[7] Pevzner, P.A., Mulyukov, Z., Dancik, V., Tang, C.L., Efficiency of database search for identification of mutated and modified proteins via mass spectrometry. Genome Res. 2001, 11, 290–299.

[8] Havilio, M., Wool, A., Large-scale unrestricted identification of post-translation modifications using tandem mass spectrometry. Anal. Chem. 2007, 79, 1362–1368.

[9] Baumgartner, C., Rejtar, T., Kullolli, M., Akella, L.M., Karger, B.L., SeMoP: A new computational strategy for the unrestricted search for modified peptides using LC-MS/MS data. J. Proteome Res. 2008, 7, 4199–4208.

[10] Searle, B.C., Dasari, S., Wilmarth, P.A., Turner, M., et al., Identification of protein modifications using MS/MS de novo sequencing and the OpenSea alignment algorithm. J. Proteome Res. 2005, 4, 546–554.

[11] Han, Y., Ma, B., Zhang, K., in:, Proc. - 2004 IEEE Comput. Syst. Bioinforma. Conf. CSB 2004, IEEE Computer Society, 2004, pp. 206–215.

[12] Chalkley, R.J., Baker, P.R., Medzihardszky, K.F., Lynn, A.J., Burlingame, A.L., In-depth analysis of tandem mass spectrometry data from disparate instrument types. Mol. Cell. Proteomics 2008, 7, 2386–2398.

[13] Na, S., Paek, E., Prediction of novel modifications by unrestrictive search of tandem mass spectra. J. Proteome Res. 2009, 8, 4418–4427.

[14] Dasari, S., Chambers, M.C., Slebos, R.J., Zimmerman, L.J., et al., TagRecon: High-throughput mutation identification through sequence tagging. J. Proteome Res. 2010, 9, 1716–1726.

[15] Na, S., Bandeira, N., Paek, E., Fast multi-blind modification search through tandem mass spectrometry. Mol. Cell. Proteomics 2012, 11.

[16] Chick, J.M., Kolippakkam, D., Nusinow, D.P., Zhai, B., et al., A mass-tolerant database search identifies a large proportion of unassigned spectra in shotgun proteomics as modified peptides. Nat. Biotechnol. 2015, 33, 743–749.

[17] Kong, A.T., Leprevost, F. V., Avtonomov, D.M., Mellacheruvu, D., Nesvizhskii, A.I., MSFragger: Ultrafast and comprehensive peptide identification in mass spectrometry-based proteomics. Nat. Methods 2017, 14, 513–520.

[18] Chi, H., Liu, C., Yang, H., Zeng, W.F., et al., Comprehensive identification of peptides in tandem mass spectra using an efficient open search engine. Nat. Biotechnol. 2018, 36, 1059–1066.

[19] Devabhaktuni, A., Lin, S., Zhang, L., Swaminathan, K., et al., TagGraph reveals vast protein modification landscapes from large tandem mass spectrometry datasets. Nat. Biotechnol. 2019, 37, 469–479.

[20] An, Z., Zhai, L., Ying, W., Qian, X., et al., PTMiner: Localization and quality control of protein modifications detected in an open search and its application to comprehensive post-translational modification characterization in human proteome. Mol. Cell. Proteomics 2019, 18, 391–405.

[21] Avtonomov, D.M., Kong, A., Nesvizhskii, A.I., DeltaMass: Automated Detection and Visualization of Mass Shifts in Proteomic Open-Search Results. J. Proteome Res. 2019, 18, 715–720.

[22] Creasy, D.M., Cottrell, J.S., Unimod: Protein modifications for mass spectrometry. Proteomics 2004, 4, 1534–1536.

[23] Geiszler, D.J., Kong, A.T., Avtonomov, D.M., Yu, F., et al., PTM-Shepherd: Analysis and Summarization of Post-Translational and Chemical Modifications From Open Search Results. Mol. Cell. Proteomics 2021, 20, 100018.

[24] Bubis, J.A., Levitsky, L.I., Ivanov, M. V., Gorshkov, M. V., Validation of Peptide Identification Results in Proteomics Using Amino Acid Counting. Proteomics 2018, 18.

[25] Stepath, M., Zülch, B., Maghnouj, A., Schork, K., et al., Systematic Comparison of Label-Free, SILAC, and TMT Techniques to Study Early Adaption toward Inhibition of EGFR Signaling in the Colorectal Cancer Cell Line DiFi. J. Proteome Res. 2020, 19, 926–937.

[26] Saei, A.A., Beusch, C.M., Chernobrovkin, A., Sabatier, P., et al., ProTargetMiner as a proteome signature library of anticancer molecules for functional discovery. Nat. Commun. 2019, 10.

[27] Narimatsu, Y., Joshi, H.J., Schjoldager, K.T., Hintze, J., et al., Exploring regulation of protein O-glycosylation in isogenic human HEK293 cells by differential O-glycoproteomics. Mol. Cell. Proteomics 2019, 18, 1396–1409.

[28] Yang, W., Jackson, B., Zhang, H., Identification of glycoproteins associated with HIV latently infected cells using quantitative glycoproteomics. Proteomics 2016, 16, 1872–1880.

[29] Qin, H., Cheng, K., Zhu, J., Mao, J., et al., Proteomics Analysis of O-GalNAc Glycosylation in Human Serum by an Integrated Strategy. Anal. Chem. 2017, 89, 1469–1476.

[30] Hansen, B.K., Gupta, R., Baldus, L., Lyon, D., et al., Analysis of human acetylation stoichiometry defines mechanistic constraints on protein regulation. Nat. Commun. 2019, 10.

[31] Musiani, D., Bok, J., Massignani, E., Wu, L., et al., Proteomics profiling of arginine methylation defines PRMT5 substrate specificity. Sci. Signal. 2019, 12, 8388.

[32] Vasaikar, S., Huang, C., Wang, X., Petyuk, V.A., et al., Proteogenomic Analysis of Human Colon Cancer Reveals New Therapeutic Opportunities. Cell 2019, 177, 1035–1049.e19.

[33] Emadali, A., Gallagher-Gambarelli, M., Quantitative proteomics by SILAC: practicalities and perspectives for an evolving approach. Medecine/Sciences 2009, 25, 835–842.

[34] Thompson, A., Schäfer, J., Kuhn, K., Kienle, S., et al., Tandem mass tags: A novel quantification strategy for comparative analysis of complex protein mixtures by MS/MS. Anal. Chem. 2003, 75, 1895–1904.

[35] Paul Zolg, D., Wilhelm, M., Schmidt, T., Médard, G., et al., Proteometools: Systematic characterization of 21 post-translational protein modifications by liquid chromatography tandem mass spectrometry (lc-ms/ms) using synthetic peptides. Mol. Cell. Proteomics 2018, 17, 1850–1863.

[36] Goloborodko, A.A., Levitsky, L.I., Ivanov, M. V., Gorshkov, M. V., Pyteomics—a Python Framework for Exploratory Data Analysis and Rapid Software Prototyping in Proteomics. J. Am. Soc. Mass Spectrom. 2013, 24, 301–304.

[37] Levitsky, L.I., Klein, J.A., Ivanov, M. V., Gorshkov, M. V., Pyteomics 4.0: Five Years of Development of a Python Proteomics Framework. J. Proteome Res. 2019, 18, 709–714.

[38] Ivanov, M.V., Levitsky, L.I., Bubis, J.A., Gorshkov, M.V., Scavager: A Versatile Postsearch Validation Algorithm for Shotgun Proteomics Based on Gradient Boosting. Proteomics 2019, 19.

[39] Szklarczyk, D., Franceschini, A., Wyder, S., Forslund, K., et al., STRING v10: protein–protein interaction networks, integrated over the tree of life. Nucleic Acids Res. 2015, 43, D447–D452.

[40] Ester, M., Kriegel, H.-P., Sander, J., Xu, X., in:, Proc. 2nd Int. Conf. Knowl. Discov. Data Min., vol. 96, 1996, pp. 226–231.

[41] Schubert, E., Sander, J., Ester, M., Kriegel, H.P., Xu, X., DBSCAN revisited, revisited: Why and how you should (still) use DBSCAN. ACM Trans. Database Syst. 2017, 42, 1–21.

[42] Pedregosa, F., Varoquaux, G., Gramfort, A., Michel, V., et al., Scikit-learn: Machine learning in Python. J. Mach. Learn. Res. 2011, 12, 2825–2830.

[43] Gorshkov, M. V., Good, D.M., Lyutvinskiy, Y., Yang, H., Zubarev, R.A., Calibration function for the orbitrap FTMS accounting for the space charge effect. J. Am. Soc. Mass Spectrom. 2010, 21, 1846–1851.

[44] Levitsky, L.I., Ivanov, M.V., Lobas, A.A., Bubis, J.A., et al., IdentiPy: An Extensible Search Engine for Protein Identification in Shotgun Proteomics. J. Proteome Res. 2018, 17.

[45] Kuznetsova, K.G., Levitsky, L.I., Pyatnitskiy, M.A., Ilina, I.Y., et al., Cysteine alkylation methods in shotgun proteomics and their possible effects on methionine residues. J. Proteomics 2021, 231, 104022.

[46] Solovyeva, E.M., Moshkovskii, S.A., Gorshkov, M. V., Identification-Free Control over the Precursor Isotopic Mass Misassignment in Orbitrap-Based Proteomics. J. Am. Soc. Mass Spectrom. 2021, 32, 218–224.

[47] Onisko, B.C., The Hydroxyproline Proteome of HeLa Cells with Emphasis on the Active Sites of Protein Disulfide Isomerases. J. Proteome Res. 2020, 19, 756–768.

[48] Shoulders, M.D., Raines, R.T., Collagen Structure and Stability. Annu. Rev. Biochem. 2009, 78, 929–958.

[49] Gorres, K.L., Raines, R.T., Prolyl 4-hydroxylase. Crit. Rev. Biochem. Mol. Biol. 2010, 45, 106–124.

[50] Afjehi-Sadat, L., Garcia, B.A., Comprehending dynamic protein methylation with mass spectrometry. Curr. Opin. Chem. Biol. 2013, 17, 12–19.

[51] Levitsky, L.I., Ivanov, M. V., Lobas, A.A., Gorshkov, M. V., Unbiased False Discovery Rate Estimation for Shotgun Proteomics Based on the Target-Decoy Approach. J. Proteome Res. 2017, 16, 393–397.

