## Supplementary Figures 1-14 for "AA_stat: intelligent profiling of *in vivo* and *in vitro* modifications from open search results"

*Journal of Proteome Research - Research Article*

**Table of contents**

**Appendix A.** Step-by-step description of the analysis of proline hydroxylation in proteins other than collagens with MSFragger/AA_stat workflow.
**S-Figure 1.** Example of mass calibration using DBSCAN clustering compensating partial mass shifts in an experimental run.
**S-Figure 2A.** Algorithm for recommendation of fixed modifications.
**S-Figure 2B.** Algorithm for recommendation of variable modifications.
**S-Figure 3.** Complex sample preparation with PTM enrichment can result in the peptide fraction containing completely oxidized Met residues.
**S-Figure 4.** The number of peptides in the mass shift of +128 Da decreases with increasing the number of allowed missed cleavages.
**S-Figure 5.** Logic for manual determination of the allowed number of modifications per peptide and/or per residue from the AA_stat HTML report.
**S-Figure 6**. Numbers of peptide identifications in open and closed searches.
**S-Figure 7.** GO analysis revealed enrichments of collagen trimer and endoplasmic reticulum (ER) components where proline hydroxylation takes place.
**S-Figure 8.** Mass spectra for peptides from protein disulfide-isomerases and thioredoxin domain-containing protein demonstrating proline oxidation.
**S-Figure 9.** Mass spectra for peptides from human histones demonstrating proline oxidation.
**S-Figure 10**. Numbers of modifications per peptide (left) and fraction of modified peptides (right) in the datasets with PTM enrichments and isotopic labeling: modeling high complex data for open search strategies.

**S-Figure 11.** Example of matching peptide sequences that differ by N-terminal Lys (KTLPSGcamCFNTPSIEKP and TLPSGcamCFNTPSIEKP) to the same experimental spectrum in open search.

**S-Figure 12.** Estimation of AA_stat localization accuracy on synthetic peptides with different modifications.

**S-Figure 13.** Mass shift overlap of AA_stat and PTM-Shepherd for 100 colon cancer samples from CPTAC S037 study.

**S-Figure 14.** Correlation of peptide numbers in overlapping mass shifts for AA_stat and for PTM-Shepherd.

**S-Tables 1-6** are provided separately as an .xlsx file.

**Appendix A.** Step-by-step description of proline hydroxylation analysis in proteins other than collagens with MSFragger/AA_stat workflow.

1. Download raw mass spectrometry data for CPTAC colon cancer samples (*https://cptac-data-portal.georgetown.edu/study-summary/S037*).
2. Download protein database in FASTA format (e.g. from *ftp://ftp.uniprot.org/pub/databases/uniprot/current_release/knowledgebase/taxonomic_divisions/*)
3. Prepare target_decoy database using Pyteomics (*https://pyteomics.readthedocs.io/en/latest/api/fasta.html#pyteomics.fasta.write_decoy_db*)
4. Prepare files with open search parameters (example is at *https://github.com/Nesvilab/MSFragger/blob/master/parameter_files/open_fragger.params*):
   Ensure precursor tolerance covers the mass range of modifications of interest; fragment tolerances, enzyme specificity, fragment_ion_series are correct. Switch of the variable and fixed modifications. The remaining can be left at default.
5. Prepare file with AA_stat parameters (*https://github.com/SimpleNumber/aa_stat/blob/master/AA_stat/example.cfg*):
   Ensure cleavage rule, fragment tolerance, enzyme specificity, ion types (fragment_ion_series) are the same as for open search. Specify a maximum number of variable modifications to report. The remaining can be left at default.
6. Convert raw mass spectrometry data to a standard format accepted by MSFragger engine:
   ```
   msconvert data.RAW --mzML --filter "peakPicking true 1-" --
   filter MS2Deisotope
   ```
   (*http://proteowizard.sourceforge.net/tools.shtml*)
7. Run AA_search (*https://github.com/SimpleNumber/aa_stat*)
   ```
   AA_search --params path_to_aastat_params \
   --MSFragger /home/MSFragger/MSFragger-2.3/MSFragger-2.3.jar\
   --dir aa_search_dir/ --mzml path_to/*.mzML \
   --os path_to_os_params
   ```
8. Open `~/aa_search_dir/report.html` in the folder with AA_search results to review a summary of data analysis. Go to the last step of open search (by clicking the "Next step" link at the top of report until it is no longer shown). Find the mass shift corresponding to the modification of interest (e.g. +15.9950 for oxidation) in the table titled as **Amino acid statistics**, "***mass shift***" column (*https://levitsky.github.io/aa_stat_report_example/*). Click on the number of peptides (in the "***# peptides in bin***" column) to see the peptides and corresponding mass shift localizations. For example, +15.9950 Da has a preferable localization at Met; however, there are other locations.

9. To programmatically evaluate the PSMs with modification of interest (e.g. proline oxidation), retrieve +15.9950.csv file from the folder ~/aa_search/os_step_n, where n is the last step of open search.
10. Group peptides by proteins they belong to and analyze for the repeated sequence motifs. If the peptide population is large, use any tool for determination of sequence motif enrichments (e.g. *http://slim.icr.ac.uk/pssmsearch/*). Run GO analysis for fast determination of protein families and/or related subunits of protein complexes (e.g. STRING, *https://string-db.org/cgi/input?sessionId=b1A4hyH6GZXd&input_page_active_form=multiple_identifiers*).
11. Compare mass shift localization with PTM annotations in publicly available PTM databases and literature (https://www.uniprot.org/uniprot/?query=*&fil=organism%3A%22Homo+sapiens+%28Human%29+%5B9606%5D%22+AND+reviewed%3Ayes, https://research.bioinformatics.udel.edu/iptmnet/ or any other source). Define a group of peptides with unannotated modification sites.
12. Visualize the annotated mass spectra for manual review of the fragments proving PTM localization. Use any of the tools.
    (e.g.
    https://pyteomics.readthedocs.io/en/latest/api/pylab_aux.html#pyteomics.pylab_aux.annotate_spectrum,
    https://spectrumviewer.org/,
    https://github.com/wenbostar/PDV,
    http://www.interactivepeptidespectralannotator.com/PeptideAnnotator.html).
13. Recompose sample population or increase the size of the sample cohort to reproduce and to cross-validate the results (e.g. 100 CPTAC colon cancer samples were analyzed).

**S-Figure 1.** Example of mass calibration using DBSCAN clustering compensating partial mass shifts in an experimental run: (A) before calibration, (B) after calibration. First sample fraction in dataset #3 has a systematic mass shift taking place from 55th to 80th minutes of the gradient time during the experimental run. If not accounted, it results in a series of uninterpretable mass shifts (0.028 Da, 15.99+0.028 Da) in the results of open search.

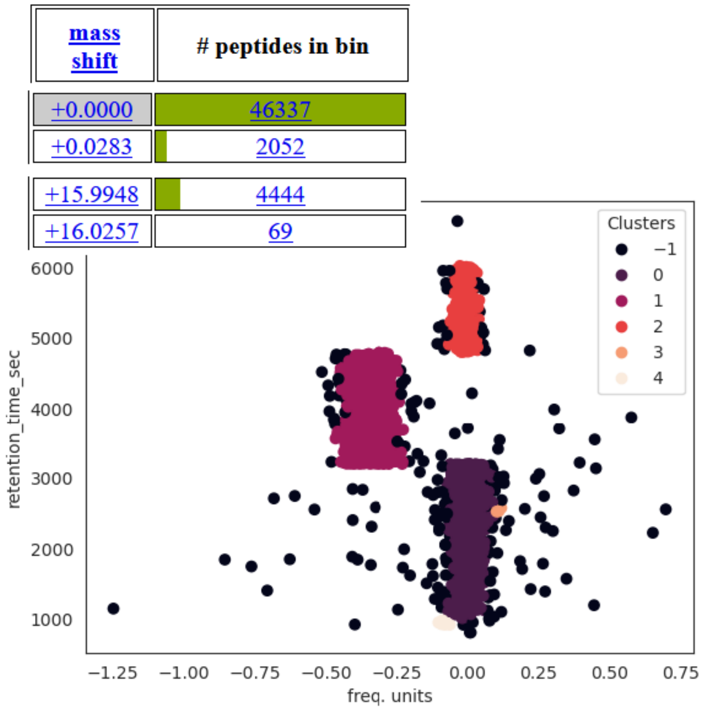


(A)

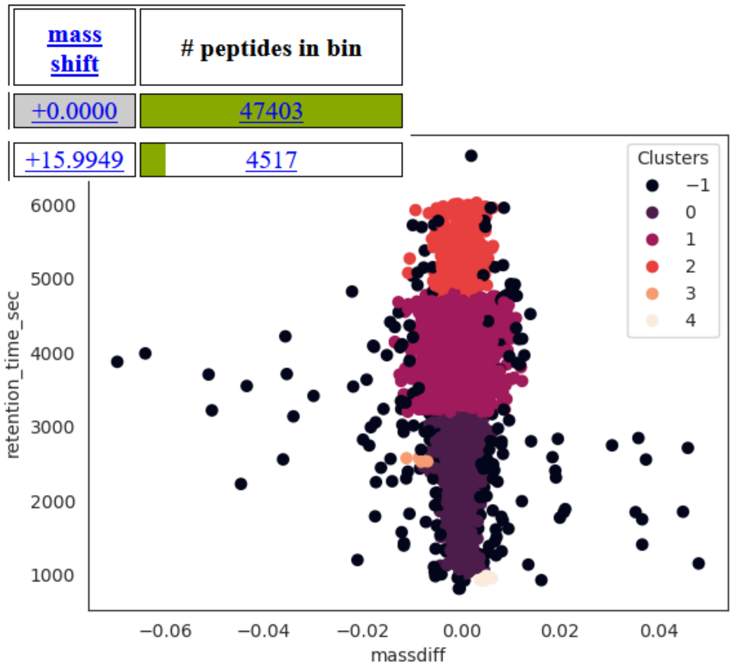


(B)

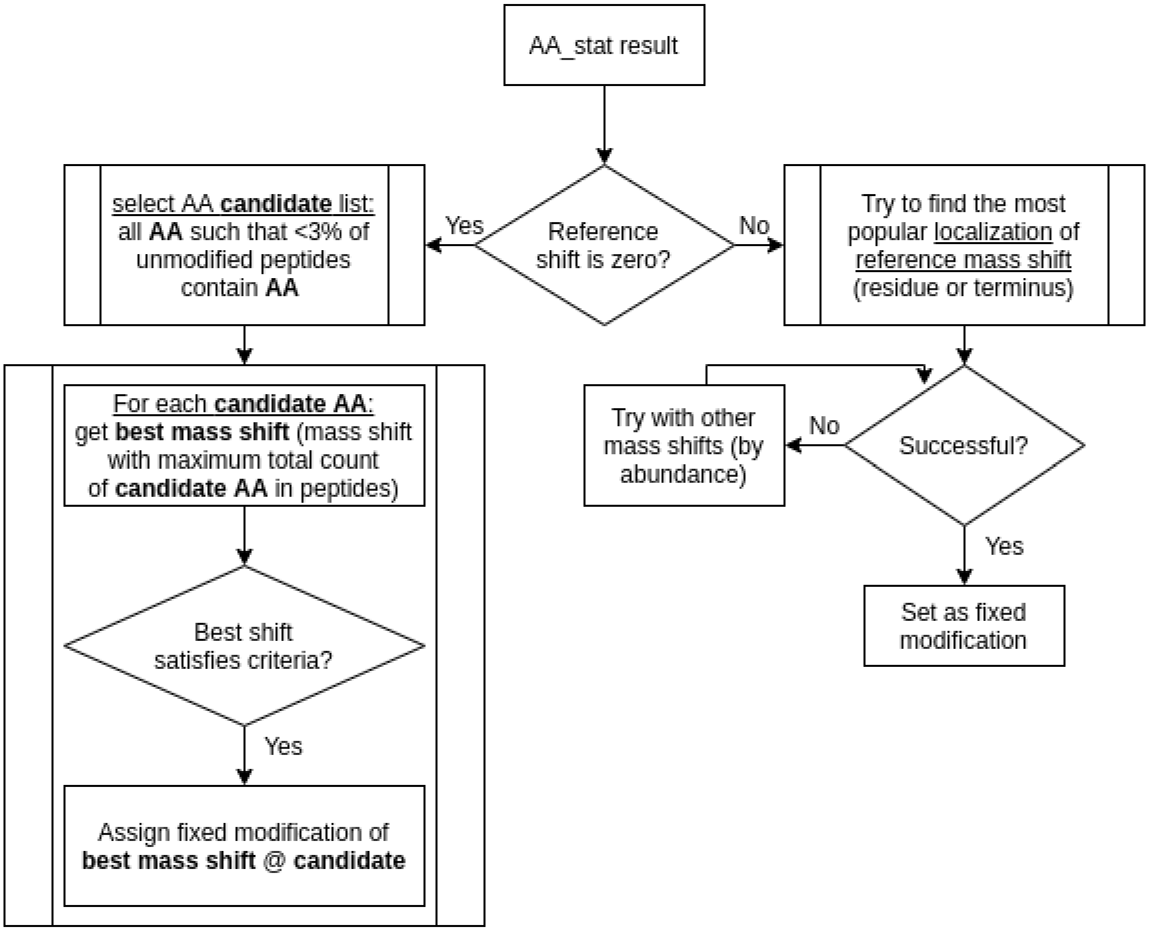


**S-Figure 2A.** Algorithm for recommendation of fixed modifications.

Briefly, the algorithm starts with checking the reference mass shift determined by AA_stat. If the reference mass shift is 0.0 Da, a list of amino acid candidates bearing fixed modifications is composed of amino acids whose occurrence at zero shift is below a threshold. Then, for each residue candidate, the mass shift containing the maximum total peptide percentage is determined. If the peptide percentage for this mass shift satisfies the selection criterion, the fixed modification is recommended. If the reference shift is not zero, preferable mass shift localizations in the non-zero reference shift are ranked. If the most preferable localization site is determined, then this mass shift is recommended as fixed modification on the preferable site. Otherwise, the procedure is repeated for the mass shift next by abundance.

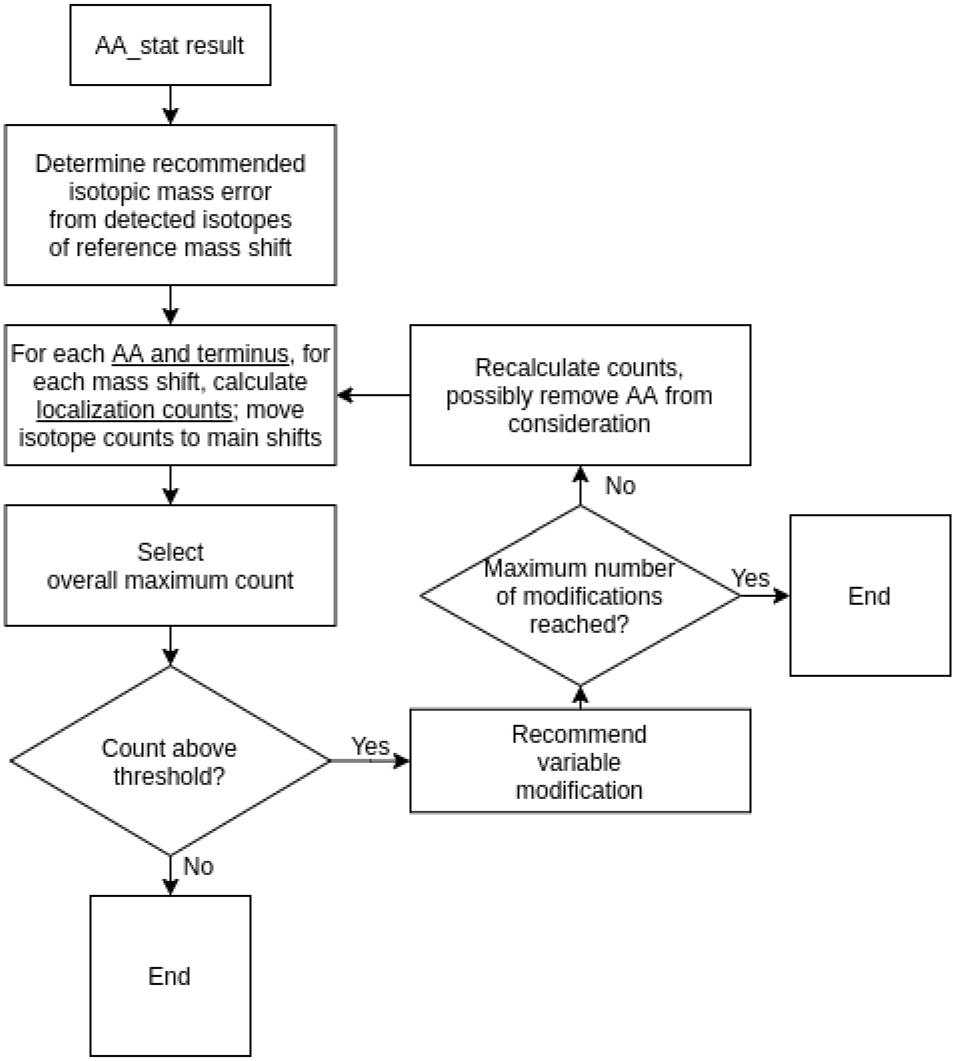


**S-Figure 2B.** Algorithm for recommendation of variable modifications. Algorithm starts with estimation of isotopic error rate in the data, and then it calculates localization counts for each amino acid and terminus in each mass shift, taking into account isotopic error counts. After selecting an amino acid with maximum localization count and checking if this count passes a predefined threshold, the variable modification is recommended, followed by checking if the maximum number of modifications is achieved. For the next round, the algorithm recalculates the localization counts excluding the previously selected amino acid from consideration.

**S-Figure 3.** Complex sample preparation with PTM enrichment can result in the peptide fraction containing completely oxidized Met residues. It will result in AA_stat's reporting the oxMet as recommended fixed modification. We suggest to consider that as additional information on the side products of the applied sample preparation. Designations: #peptides is the number of peptides with Met residues identified in close search with Met oxidation as variable modification, #oxMet is the number of oxidized Met residues, #Met is the number of non-oxidized Met residues, mrate is the amino acid modification rate, mrate = #oxMet/(#oxMet + #Met)*100%.

| Dataset | #peptides | #oxMet | #Met | mrate, % |
|---|---|---|---|---|
| #6 N-glycosylation dataset PXD003561, subset A301 | 151 | 151 | 0 | 100.0 |
| #11 N-glycosylation dataset PXD003561, subset ACH | 397 | 396 | 3 | 99.25 |

**S-Figure 4.** The number of peptides in the mass shift of +128 Da decreases with increasing the number of allowed missed cleavages. Abundant mass shift of +128 Da can occur in open search due to underestimation of missed cleavage (mc) rates in tryptic digestion.

| dataset | #peptides in +128Da, mc = 1 | #peptides in +128Da, mc = 2 | #peptides in +128Da, mc = 3 | #peptides in +128Da, mc = 4 |
|---|---|---|---|---|
| #12 methylation dataset PXD009070, subset HF170411 | 104 | 0 | - | - |
| #13 PXD014565 SILAC subset 22200 | 240 | 75 | 0 | - |
| #14 PXD014565 SILAC subset 22204 | 245 | 68 | 0 | - |
| #15 PXD014565 SILAC subset 22208 | 243 | 79 | 55 | 0 |
| #16 CPTAC colon cancer data from study S037 (sample 01) | 85 | 0 | - | - |

Recommended, variable

| | isotope error | K | K | M | K | R | K | K |
|---|---|---|---|---|---|---|---|---|
| value | 1 | +42.0110 | +8.0140 | +15.9950 | +50.0254 | +10.0082 | +216.1344 | +58.0394 |

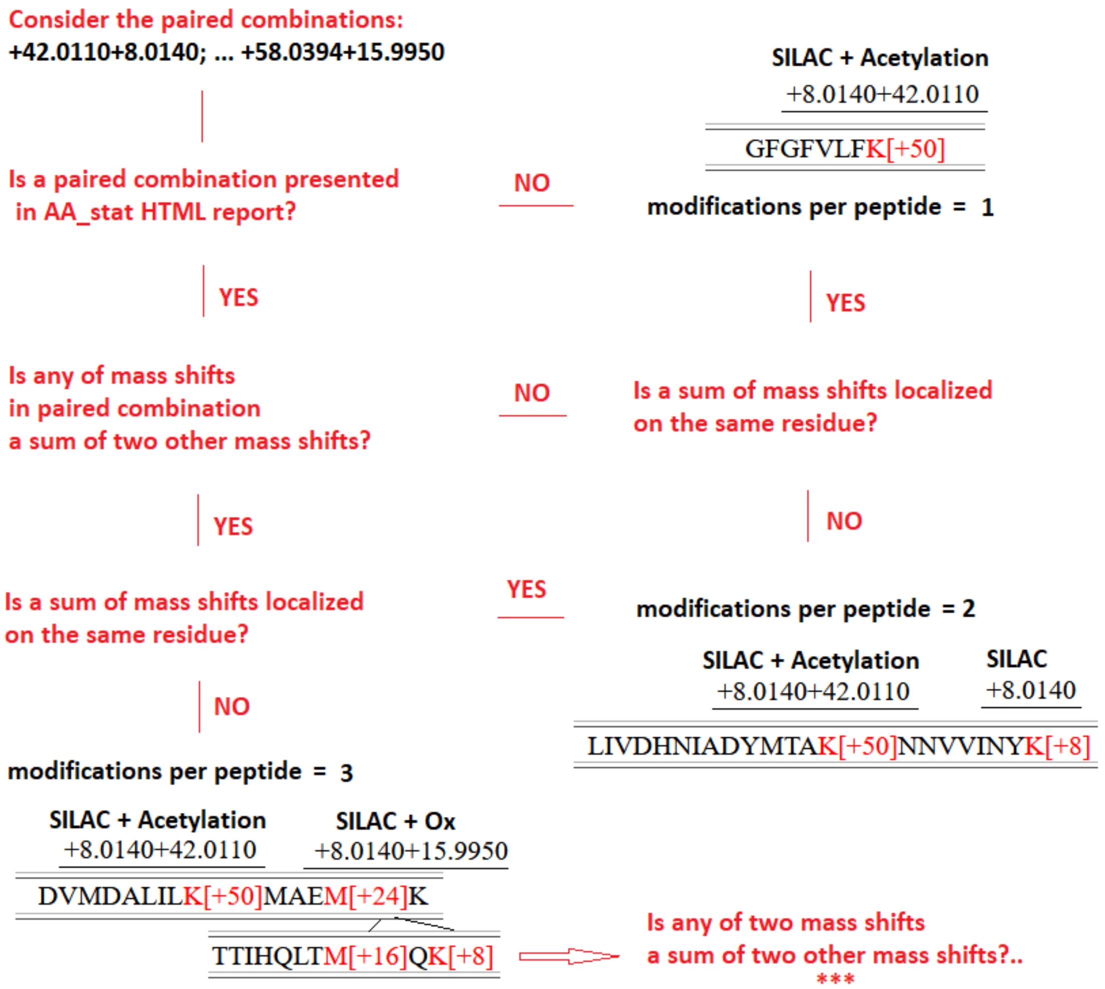


**S-Figure 5.** Logic for manual determination of the allowed number of modifications per peptide and/ or per residue from the AA_stat HTML report. The conclusion on the number of allowed modifications per peptide is based on recursive counting of paired combinations of AA_stat-recommended variable modifications, followed by counting the peptides in such combinations. If the fraction of peptides bearing specific combinations of modifications is reasonably high, then the recommended number of modifications increases. If a sum of mass shifts is localized on the same site, then the increased number of modifications per site should be taken into account. To improve identification efficiency for search engines allowing only one modification per residue, a sum of mass shifts can be specified as variable modification.

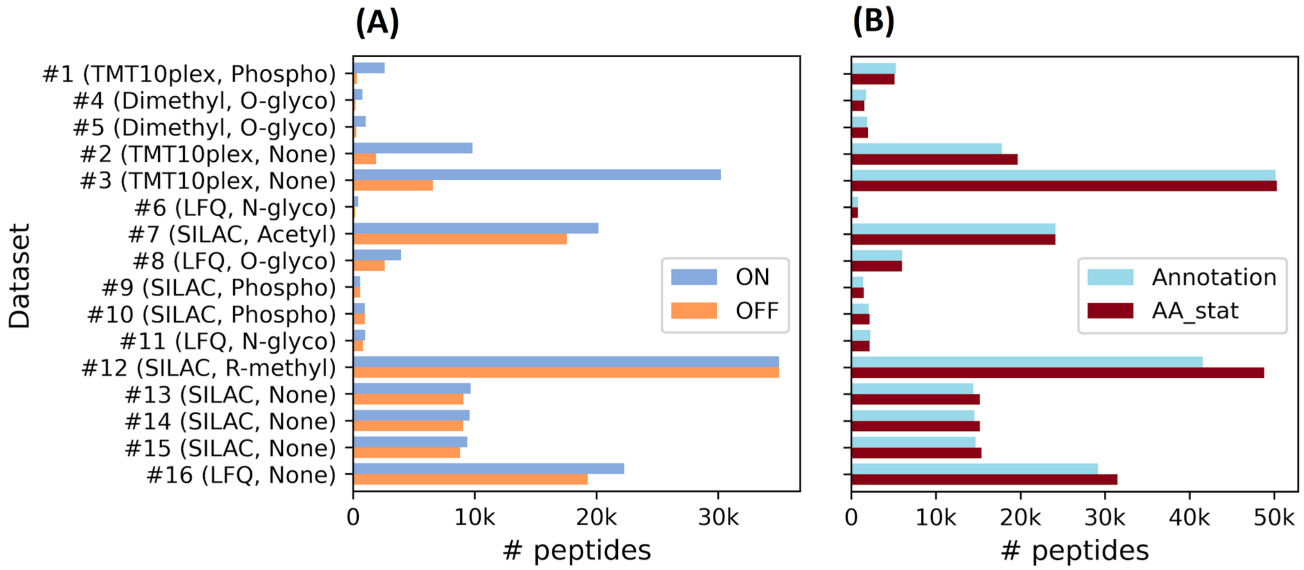


**S-Figure 6**. Numbers of peptide identifications in open and closed searches. (A): distantly located modifications significantly decrease the number of matched unmodified peptide isoforms in open search (the effect is modeled by switching ON or OF the fixed modifications and is observed especially significant for TMT and dimethyl labeling data). (B): AA_stat-retrieved top abundant mass shift localizations allow to increase the yield of closed searches. Y-axis labels correspond to subsets as listed in **S-Tables 3-4**. “None” means no enrichment. For open searches, the number of peptides is the sum of peptide counts in all mass shifts as reported by AA_stat. For closed searches, the number of peptides as reported by Scavager is shown. Search results were obtained using either the annotated (from a respective study) or AA_stat-recommended sets of modifications. To retrieve modifications, AA_stat was run with the AA_search wrapper.

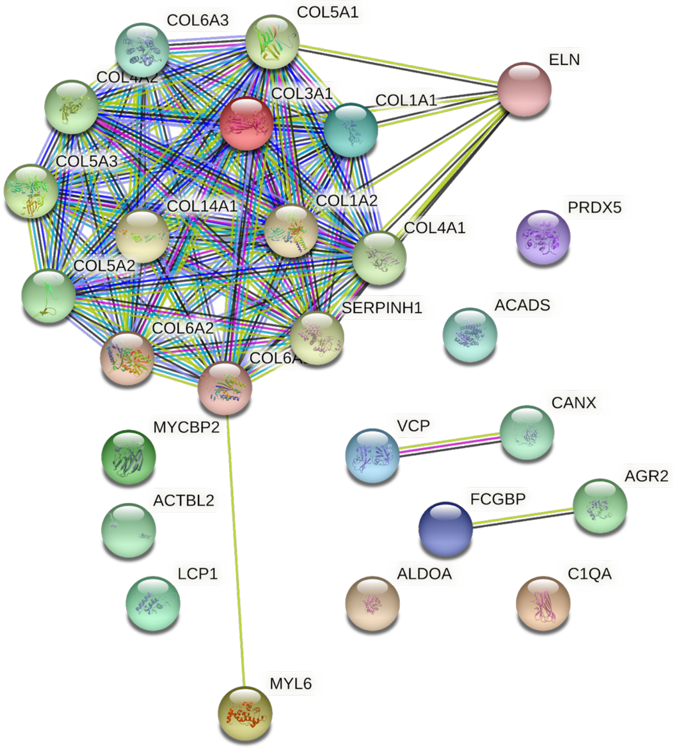


**S-Figure 7.** GO analysis reveals enrichments of collagen trimer and endoplasmic reticulum (ER) where proline hydroxylation takes place.

**S-Figure 8.** Mass spectra for peptides from protein disulfide-isomerases (PDIA1, PDIA3, PDIA4, PDIA5 and PDIA6) and thioredoxin domain-containing protein (TXNDC5) suggesting proline oxidation. Fragment tolerance 0.05 Da. Peptides identified across 100 CPTAC colon cancer samples (S037).

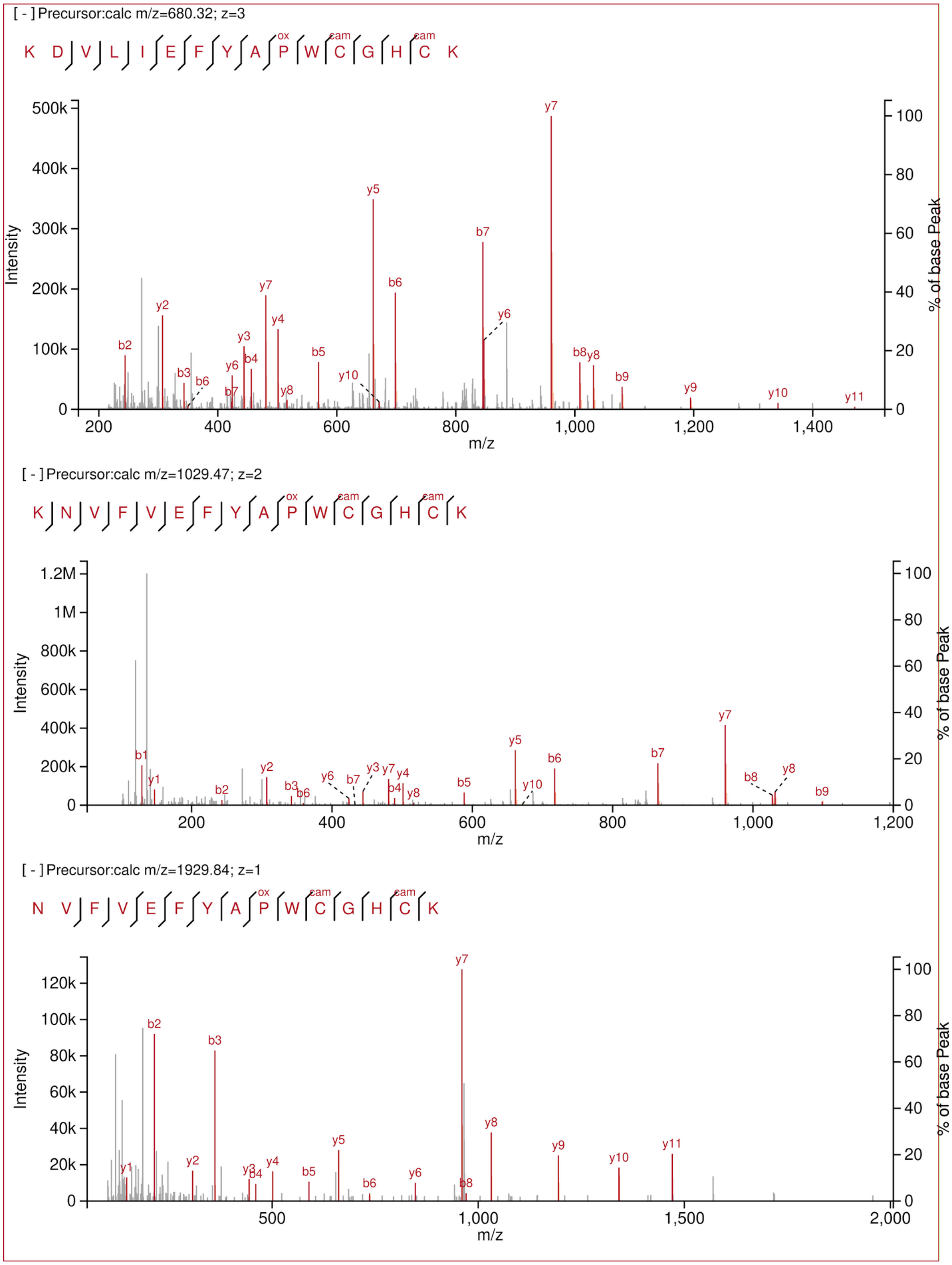

[ – ] Precursor:calc m/z=1910.86; z=1

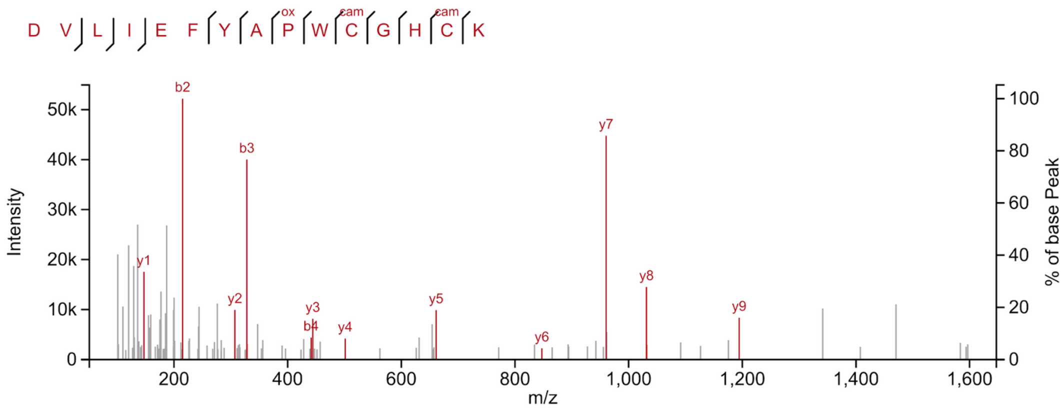


[ – ] Precursor:calc m/z=1325.56; z=1

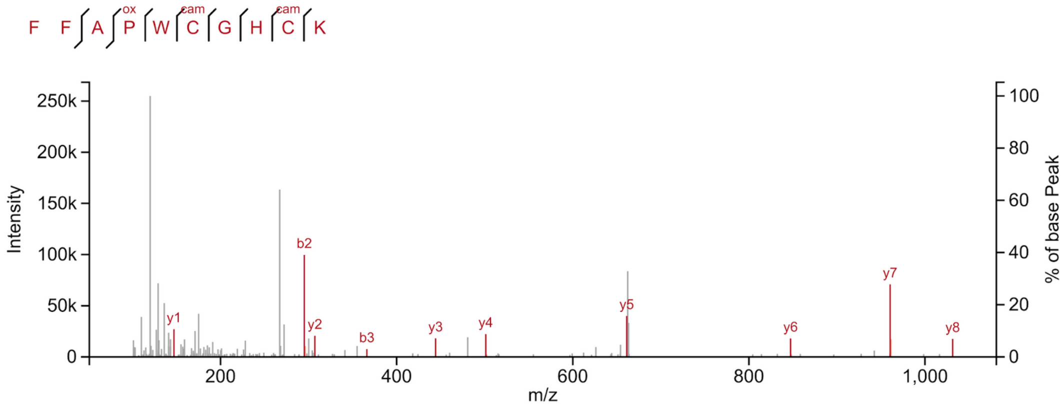


[ – ] Precursor:calc m/z=1448.66; z=2

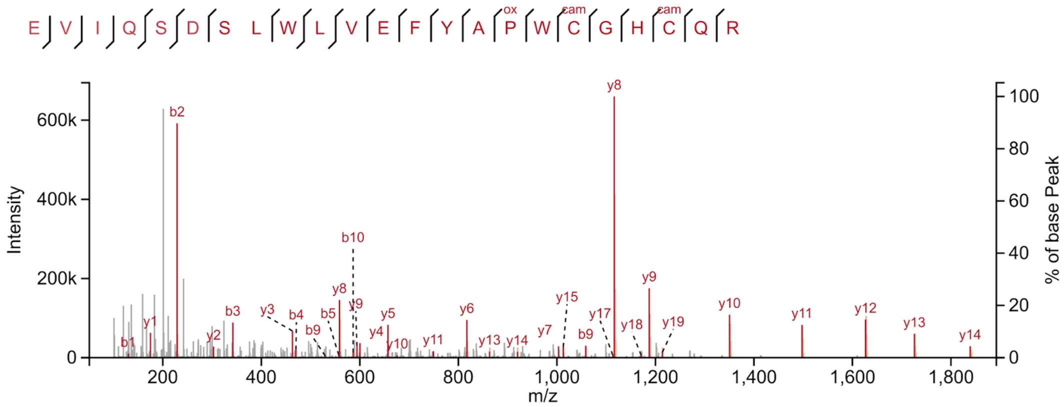

**S-Figure 9.** Mass spectra for peptides from human histones (H2B1N, H2B1O, H2BFS, H2B1C, H2B1D, H2B2F, H2B1J, H2B1K, H2B1A, H2B2E, H2B1H, H2B3B, H2B1B, H2B1L, H2B1M) demonstrating proline oxidation. Fragment tolerance 0.05 Da.

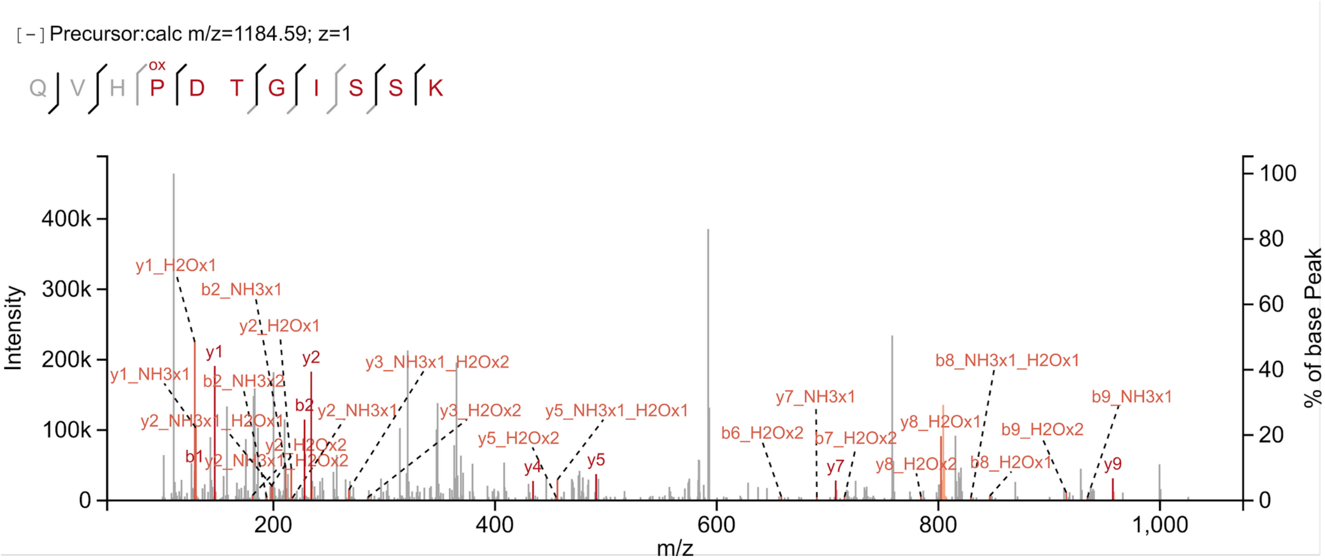

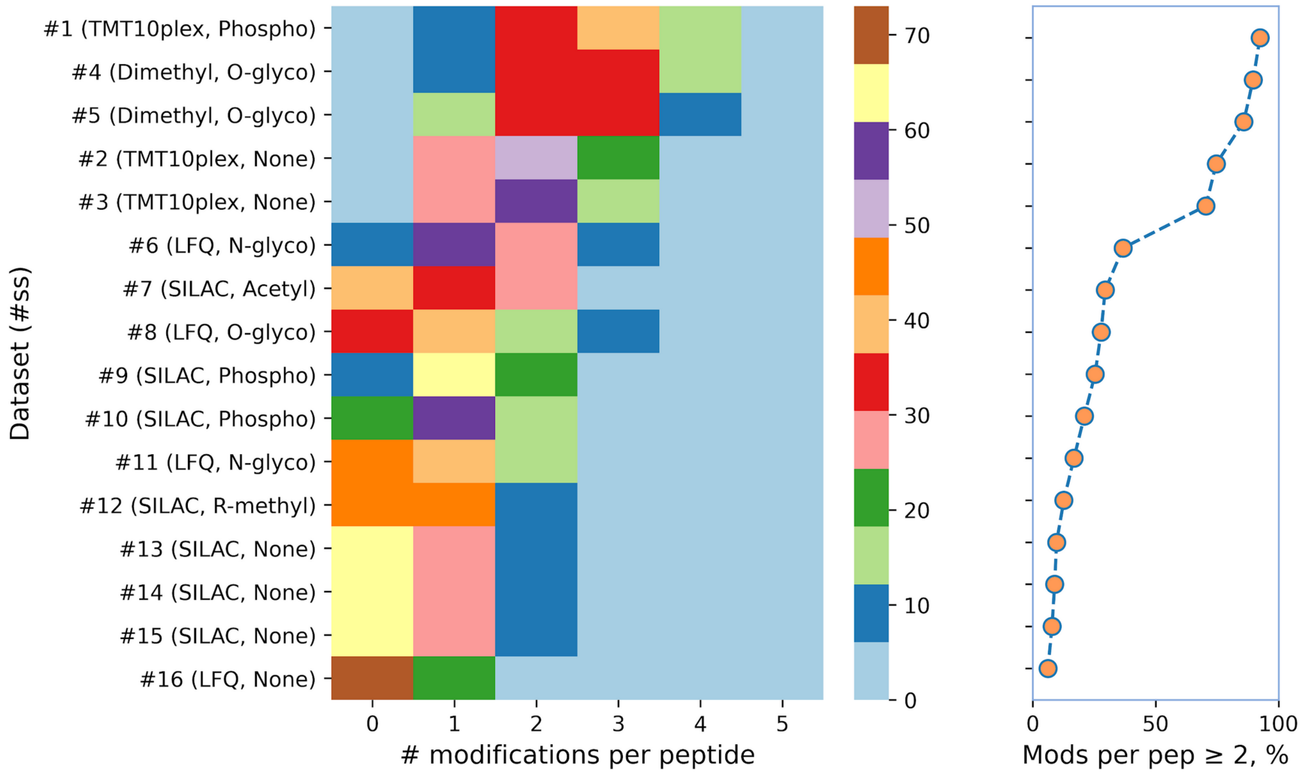


**S-Figure 10**. Numbers of modifications per peptide (left) and fraction of modified peptides (right) in the datasets with PTM enrichments and isotopic labeling: modeling high complex data for open search strategies. Color bar shows relative amounts of peptides with modifications (%) according to the data provided in **S-Table 3**. Y-axis labels correspond to #Subset (Labeling, Enrichment) as provided in **S-Table 3**. “None” means no enrichment. Numbers of modifications per peptide were calculated based on the closed search results obtained using annotated sets of fixed and variable modifications from the respective studies (**S-Table 1**).

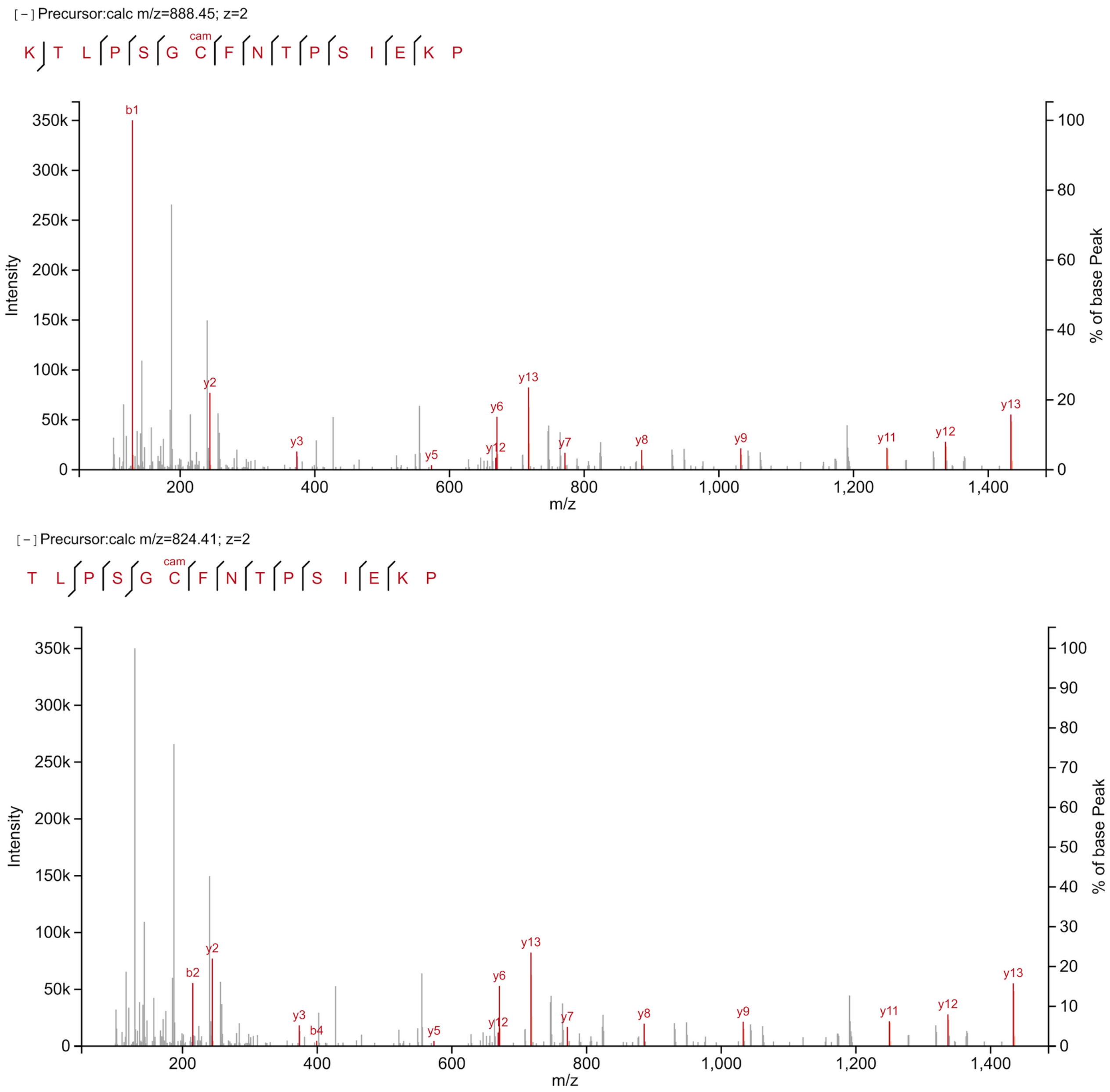


**S-Figure 11.** Example of matching peptide sequences that differ by N-terminal Lys (KTLPSGcamCFNTPSIEKP and TLPSGcamCFNTPSIEKP) to the same experimental spectra in open search. Although the b-ion series for TLPSGcamCFNTPSIEKP match two peaks, KTLPSGcamCFNTPSIEKP peptide matches one abundant b-ion that further results in a higher hyperscore accounting for intensity values.

| Modification | mass_diff, Da | # pep in set | aa_stat_indentified | aa_stat_localized | % localized |
|---|---|---|---|---|---|
| Acetyl | 42.010565 | 200 | 161 | 126 | 78 |
| Biotinyl | 226.077598 | 200 | 165 | 145 | 87 |
| Butyryl | 70.041865 | 200 | 172 | 139 | 80 |
| Citrullin | 0.984016 | 200 | 95 | 90 | 94 |
| Crotonyl | 68.026215 | 200 | 156 | 134 | 85 |
| Dimethyl | 28.0313 | 600 | 256 | 150 | 58 |
| Formyl | 27.994915 | 200 | 161 | 130 | 80 |
| Glutaryl | 114.031694 | 200 | 166 | 129 | 77 |
| GlyGlycyl | 114.042927 | 200 | 162 | 121 | 74 |
| Hydroxyisobutyryl | 86.036779 | 200 | 160 | 131 | 81 |
| Hydroxyproline | 15.994915 | 191 | 60 | 45 | 75 |
| Malonyl | 86.000394 | 200 | 147 | 135 | 91 |
| Methyl | 14.01565 | 400 | 265 | 144 | 54 |
| Nitrotyr | 44.985078 | 176 | 138 | 132 | 95 |
| Phosphorylation | 79.966331 | 176 | 146 | 128 | 87 |
| Propionyl | 56.026215 | 200 | 170 | 133 | 78 |
| Succinyl | 100.016044 | 200 | 167 | 134 | 80 |
| Trimethyl | 42.04695 | 200 | 132 | 74 | 56 |
| Unmodified | 0.0 | 688 | 478 | 478 | 100 |

**S-Figure 12.** Estimation of AA_stat localization accuracy on synthetic peptides with different modifications. *# pep in set* - actual number of synthesized peptides; *aa_stat_identified* - number of peptides found in open search; *aa_stat_localized* - number of peptides correctly localized by AA_stat; and *% localized* - percentage of correctly localized identified peptides.

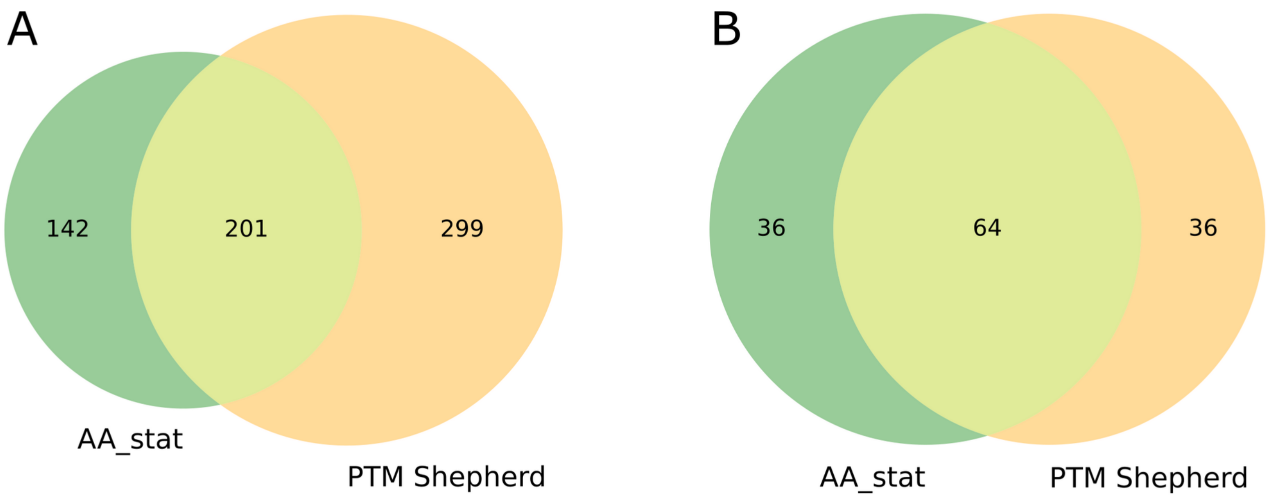


**S-Figure 13.** Mass shift overlap of AA_stat and PTM-Shepherd for 100 colon cancer samples from CPTAC S037 study: (**A**) all detected mass shifts; and (**B**) top 100 mass shifts by numbers of peptides (**B**). Values of mass shifts are considered equal if they differ by less than 0.01 Da.

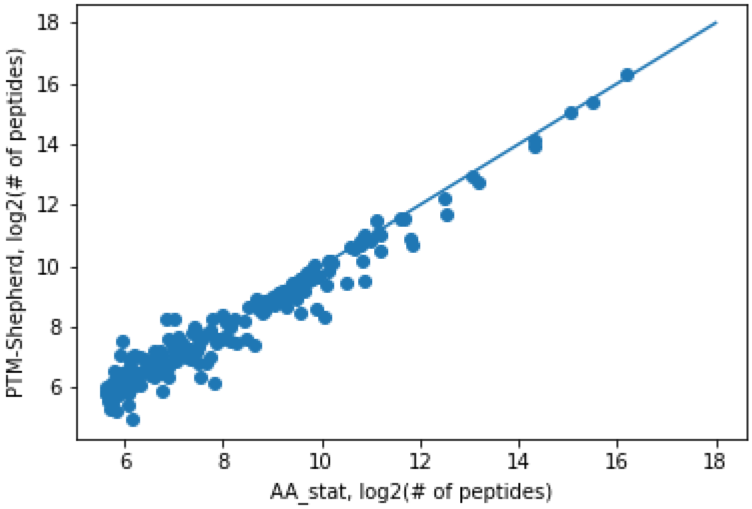


**S-Figure 14.** Correlation of peptide numbers in overlapping mass shifts for AA_stat (X axis) and for PTM-Shepherd (Y axis). All values are *log*-scaled.
